# Second-Generation Cap Analogue Prodrugs for Targeting Aberrant Eukaryotic Translation Initiation Factor 4E (eIF4E) Activity in Drug-Resistant Melanoma

**DOI:** 10.1101/2024.09.25.614990

**Authors:** Emilio L. Cárdenas, Rachel L. O’Rourke, Arya Menon, Gabriela Vega-Hernández, Jennifer Meagher, Jeanne Stuckey, Amanda L. Garner

**Affiliations:** Department of Medicinal Chemistry, College of Pharmacy, University of Michigan, Ann Arbor, Michigan 48109, United States; Program in Chemical Biology, University of Michigan, Ann Arbor, Michigan 48109, United States; Life Science Institute, University of Michigan, Ann Arbor, Michigan 48109, United States; Rogel Cancer Center, University of Michigan, Ann Arbor, Michigan 48109, United States

## Abstract

Melanoma is the deadliest form of skin cancer with a 5-year survival rate of less than 20%. While significant strides have been made in the field of kinase-targeted and immune-based therapies for melanoma, the development of resistance to these therapeutic agents has hindered the success of treatment. Drug-resistant melanoma is particularly reliant on enhanced cap-dependent translation to drive the production of oncoproteins that promote growth and survival. The m^7^GpppX cap-binding protein eukaryotic translation initiation factor 4E (eIF4E) is the rate-limiting factor of cap-dependent translation initiation, and its overexpression in melanoma tumors has been shown to drive resistance to BRAF^V600E^ kinase-targeted inhibitors. These findings point to eIF4E-targeted therapies as a promising strategy to overcome drug resistance in melanoma. Herein, we build upon our previous work of developing cell-permeable cap analogue inhibitors to design second-generation cap analogues that inhibit eIF4E-mediated cap-dependent translation in drug-resistant melanoma cells.

## INTRODUCTION

Melanoma is a cancer occurring through the uncontrolled growth of pigment-producing melanocytes found in the skin. Although melanoma is not the most common form of skin cancer, it is the deadliest with a high probability for the formation of metastatic disease which has a 5-year survival rate of only ∼20%.^1^ Indeed, while metastatic melanoma represents <1% of cases, it accounts for >80% of skin cancer-related deaths.^2^ Fortunately, the advent of targeted- and immune-based therapies over the past decade has had a profound impact on the treatment of melanoma. Standard of care for advanced metastatic disease now includes treatment with immune checkpoint inhibitors, either alone or in combination with targeted inhibitors of BRAF^V600E^ or MEK kinases.^2, 3^ Despite this remarkable progress that has been made in melanoma treatment leading to substantial improvements in survival,^3^ drug resistance to these agents has emerged as a significant impediment towards the goal of curing the majority of patients (∼80%) presenting with advanced melanoma.^2, 4, 5^ In these patients, drug resistance is primarily due either to acquired resistance to targeted therapies or innate resistance to immunotherapy.^1^ As the incidence of melanoma continues to rise disproportionally in comparison to other cancer types, there is an unmet need to develop new therapies to advance the treatment of the growing population of patients with drug-resistant, metastatic melanoma.

In melanoma, activation of both the RAS/RAF/MAPK and PI3K/AKT/mTOR pathways converge on eIF4E, the m^7^GpppX cap-binding protein, thereby promoting cap-dependent translation via phosphorylation of eIF4E and 4EBP1, respectively (Figure 1). Indeed, melanoma patients exhibit high levels of eIF4E and phospho-eIF4E and -4EBP1, each correlating with greater metastatic potential and poor patient survival.^6-8^ More recent studies have strengthened the role that eIF4E plays in the disease, demonstrating that activation of eIF4E and cap-dependent mRNA translation drives resistance to the BRAF^V600E^-targeted drug, vemurafenib,^9, 10^ and correlates with response to anti-PD-1/PD-L1 immunotherapies, both standard of care treatments for melanoma patients.^11, 12^ With respect to vemurafenib resistance, while sensitive cells and tumors showed decreased cap-dependent translation due to vemurafenib-induced stabilization of the eIF4E/4EBP1 protein-protein interaction (PPI), drug-resistant melanoma exhibited increased dependence on eIF4E and enhanced levels of the pro-translation eIF4E/eIF4G PPI (Figure 1).^9, 10^ Importantly, co-treatment with inhibitors of the eIF4E/eIF4G PPI or eIF4A was found to re-sensitize cells to vemurafenib, indicating the potential for development of eIF4F-targeted combination therapy to overcome resistance in BRAF^V600^-mutant melanomas.^9^

**Figure 1.**
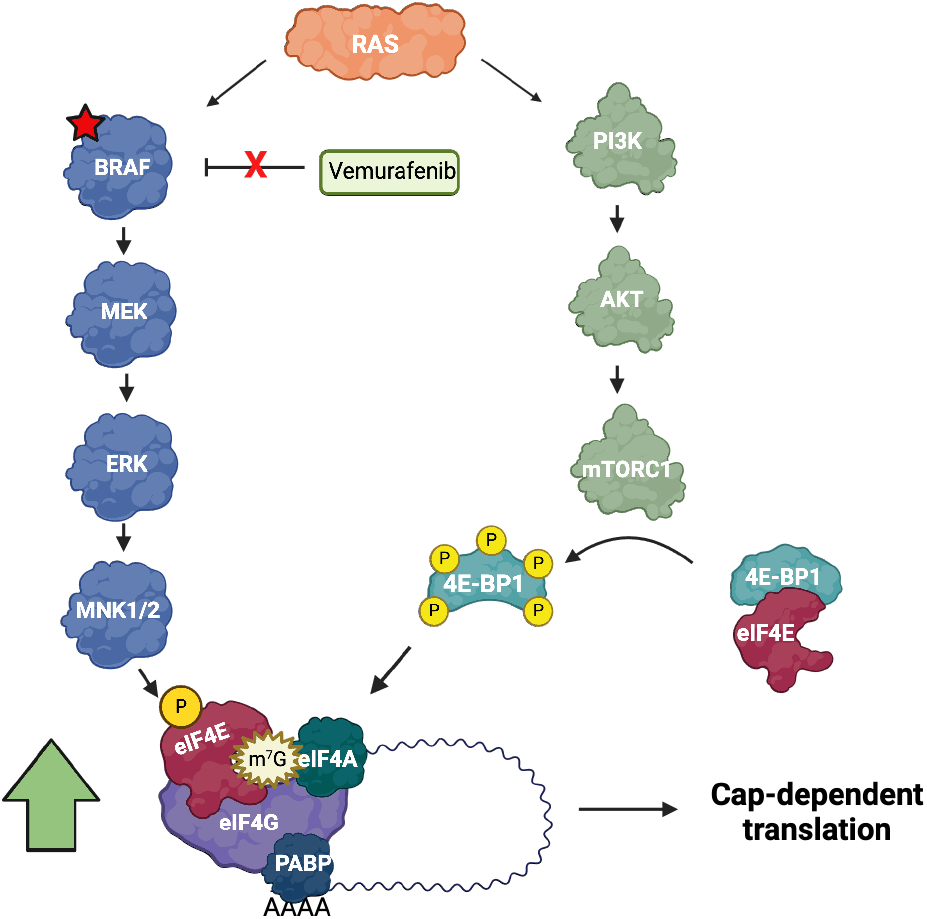
Activation of eIF4E and cap-dependent translation in vemurafenib-resistant melanoma.

Recently, our laboratory reported the design and synthesis of cell-permeable small molecule inhibitors targeting the eIF4E cap-binding site.^13^ Using a prodrug design approach inspired by the FDA-approved acyclic nucleoside phosphonate prodrugs adefovir and tenofovir dipivoxil, we reported first-generation cap analogues containing a bis-POM (*Z*)-4-phosphono-but-2-enyl linker (**1**, Figure 2).^14^ Importantly, these compounds displayed promising on-target anti-proliferative activity indicating their potential for further development to enable investigation of the therapeutic potential of directly targeting eIF4E in human cancers. Using a structure-guided approach, herein we report the design and synthesis of second-generation cap analogues as well as characterization of these inhibitors in cellular models of melanoma resulting in the uncovering of eIF4E-targeted inhibitors demonstrating inhibition of cap-dependent translation and selective anti-proliferative activity in drug-resistant melanoma cells.

**Figure 2.**
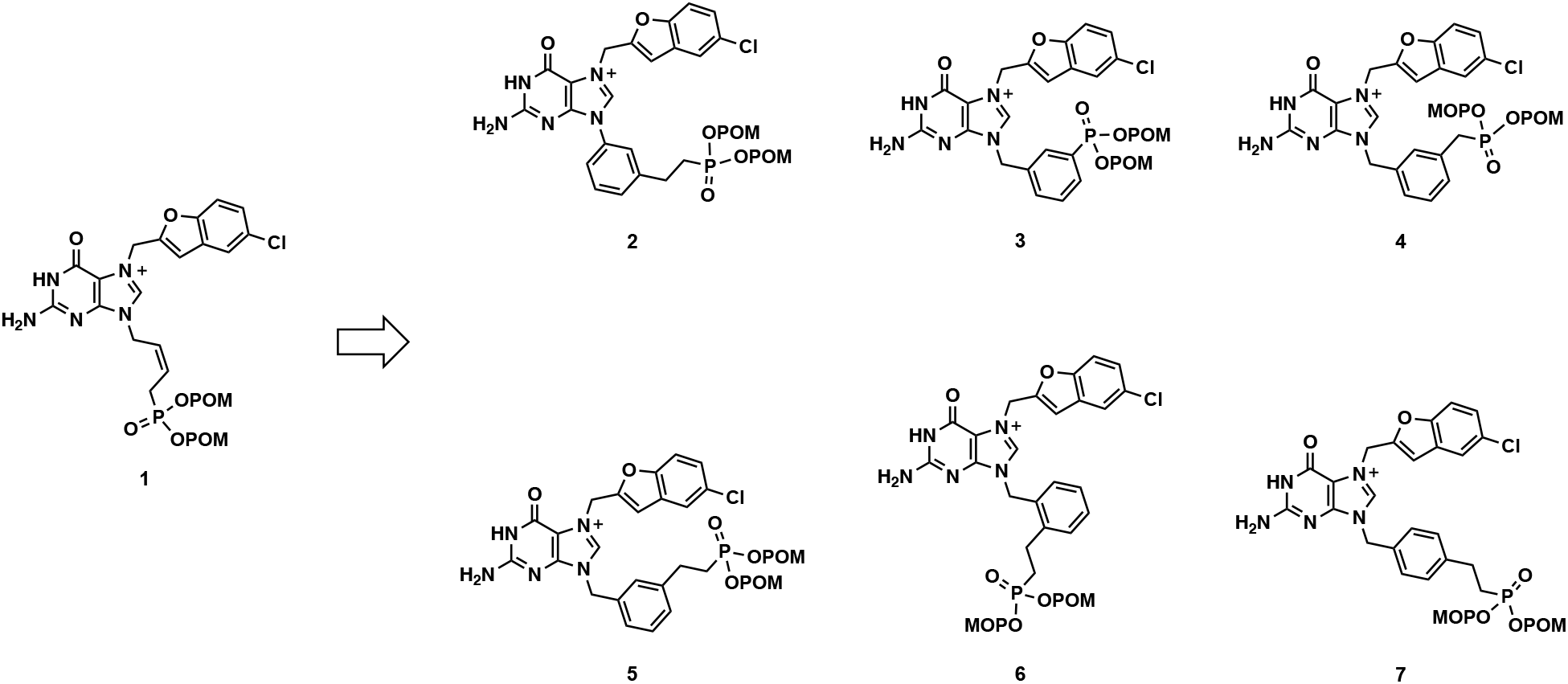
Structures of first-(**1**) and second-generation (**2−7**) nucleoside phosphonate prodrug inhibitors of eIF4E. POM = pivaloyloxymethyl.

## RESULTS AND DISCUSSION

Structural studies of cap analogue inhibitors bound to eIF4E, including our previously synthesized acyclic bis-POM cap analogues,^13, 15^ demonstrated that bioisosteric replacement of the canonical ribose sugar of m^7^GMP was plausible with an appropriate conformationally restricted linker between the nucleobase and phosphonate. Specifically, our bis-POM (*Z*)-4-phosphono-but-2-enyl linker (**1**, Figure 2) relied upon the concave shape of the cap-binding domain of eIF4E resulting in hydrogen-bonding interactions between the phosphonate and R157 located in the positively-charged cleft typically accessed by the triphosphate of the endogenous m^7^GpppX cap structure.^13^ With this structural insight in mind, we posited that an alternative *N*-9-substituted linker such as a *N*-9-phenyl-or -benzyl–substituted linker could mimic the *cis*-conformation that positioned the (*Z*)-butenyl phosphonate in proximity to the critical cationic residues that are imperative for the potency of m^7^GMP cap-based inhibitors. Indeed, others have employed similar linker design strategies, including previous work towards the generation of *N*-9-benzylguanine phosphonates that displayed extremely potent inhibition of purine nucleoside phosphorylase.^16^ Additional work demonstrated that 7-methylguanine derivatives containing *N*-9-phenyl and -benzyl linkers could serve as potent anti-viral agents through inhibition of the PB2 cap-binding domain associated with “cap-snatching” in influenza A.^17^ Based on this precedent, we synthesized a library of bis-POM cap analogues containing various *N*-9-phenyl and -benzyl linkers with a *N*-7-methyl(benzofuran) substituent due to its superior activity in our previous work (compounds **2**−**5**, Figure 2; please see the Supporting Information for synthesis details).^13^

Bis-POM cap analogue prodrugs **2**−**5** were first profiled in vemurafenib-resistant BRAF^V600E^-mutant A2058 melanoma cells to evaluate anti-proliferative activity using the CellTiter-Glo cell viability assay (Figure 3A).^9^ Following single-point analysis at 50 μM, *meta*-substituted *N*-9-benzyl analogue **5** was found to be the most potent prodrug in this series inducing near complete cell death. Closely related *N*-9-phenyl analogue **2** also showed appreciable anti-proliferative activity at this concentration. Prodrugs **3** and **4** were found to be less active, presumably due to decreased linker length separating the benzyl and phosphonate moieties which may prevent the phosphonate from interacting with key amino acid residues in the eIF4E binding pocket.^13, 15^ Subsequent dose dependence analysis with **5** showed that it exhibited an EC_50_ value of 40 μM

**Figure 3.**
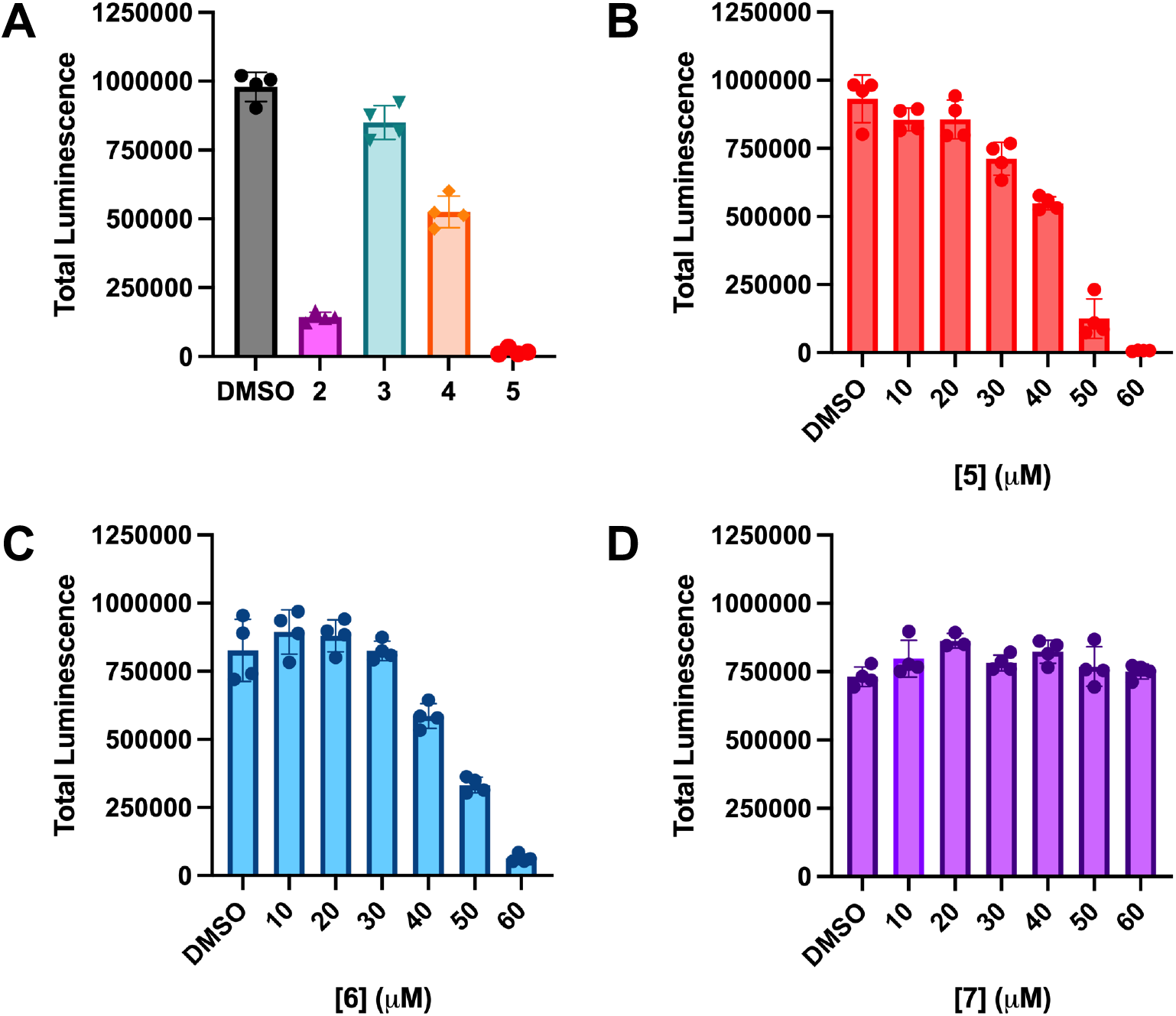
Anti-proliferative activity of second-generation bis-POM cap analogue prodrugs. (A) Single-point activity in A2058 cells (50 μM, 48 h). Dose-dependent anti-proliferative activity of (B) **5**, (C) **6**, and (D) **7**.

(Figure 3B), an activity in-line with that observed with the first-generation prodrug **1** in MiaPaca-2 cells.^13^ We then asked whether the position of the phosphonate on the benzyl linker would affect cellular activity. To explore this structure-activity relationship, we prepared and tested the corresponding *ortho*- and *para*-substituted derivatives **6** and **7** in the CellTiter-Glo assay (structures in Figure 2; please see the Supporting Information for synthesis details). While **7** was found to be inactive in this assay, likely due to improper positioning of the phosphonate group, **6** was nearly equipotent to **5** with an EC_50_ value of 51.3 μM (Figures 3C and 3D).

As bis-POM cap analogue prodrugs **5** and **6** both displayed dose-dependent anti-proliferative activity, we next confirmed their competitive inhibition of eIF4E using a fluorescence polarization (FP) assay employing fluorescein-labeled m^7^GTP as a competitive probe.^18^ For biochemical assays, phosphonic acids of **5** and **6** (**5-PA** and **6-PA**, Figure 4A) were synthesized via hydrolysis of the bis-POM phosphonate esters (please see the Supporting Information for synthesis details). In FP, **5-PA** displayed single digit micromolar competitive binding activity with an IC_50_ value of 3.5 μM, while **6-PA** was less active with an IC_50_ value of 14.7 μM (Figure 4B). As an orthogonal target engagement assay, a cellular thermal shift assay (CETSA)^19^ was performed using HeLa cell lysate to confirm engagement of **5-PA** and **6-PA** with eIF4E in the cellular environment. Stabilization of eIF4E, which indicates binding, was visualized using Western blot. In-line with the FP assay, **5-PA** showed greater stabilization of eIF4E, confirming that it binds more tightly to eIF4E (Figures 4C and 4D).

**Figure 4.**
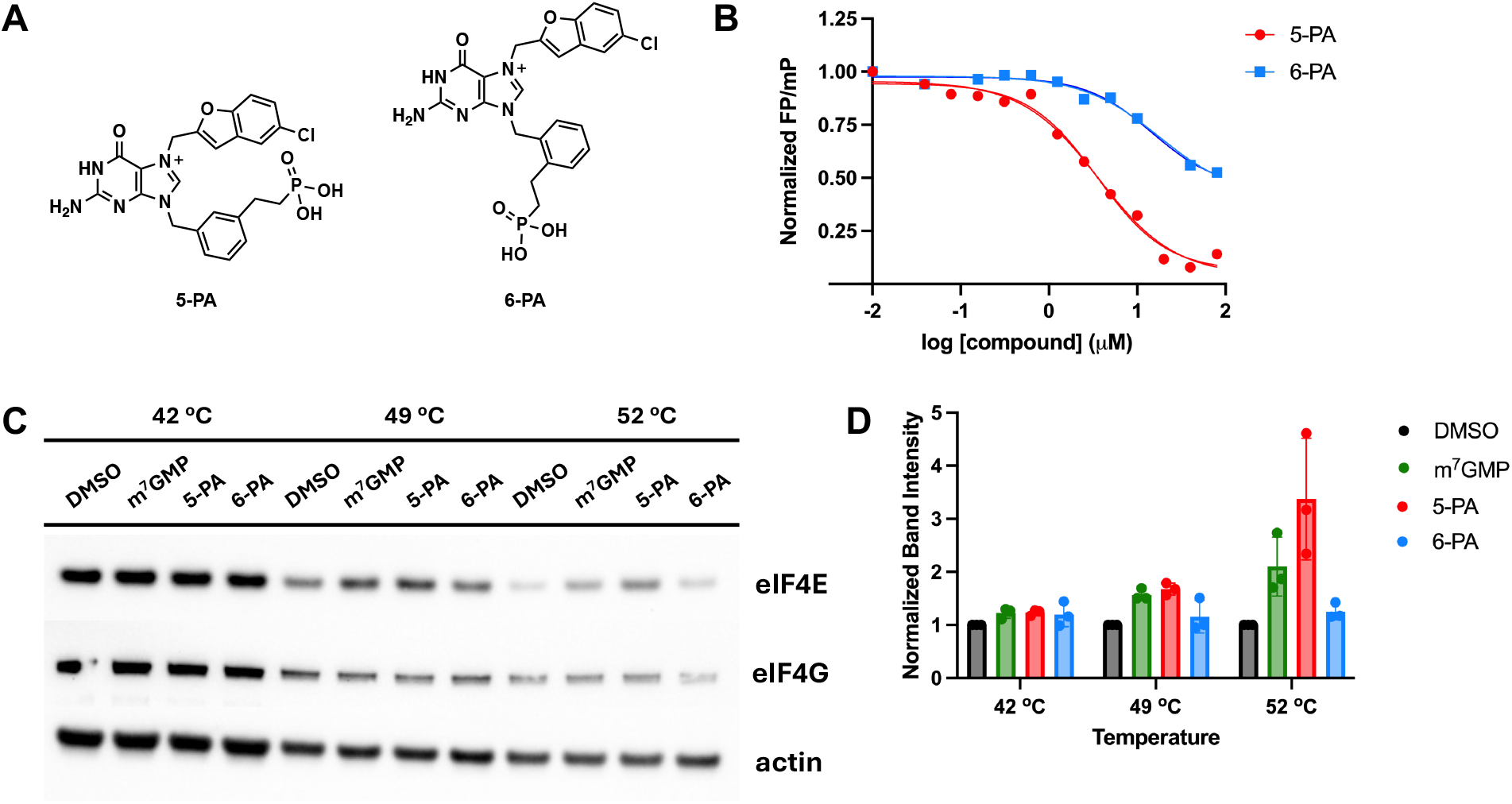
Target engagement of second-generation cap analogues. (A) Structures of phosphonic acids **5-PA** and **6-PA**. (B) Biochemical inhibitory activity measured using a FP assay. (C) Stabilization of eIF4E by **5-PA** and **6-PA** in HeLa cell lysate as determined via CETSA. Proteins were detected via Western blot. (D) Quantitation of CETSA data in comparison to DMSO as a control.

To provide structural insight into the binding mechanism of **5-PA**, an X-ray crystal structure of this compound co-crystalized in the cap-binding domain of eIF4E was resolved with a resolution of 2.1 Å resolution with a dimer in the asymmetric unit (Figure 5). The structure solved with m^7^GDP bound to one protein chain and **5-PA** occupying the cap binding site in the other protein chain. The guanine core scaffold of **5-PA** was sandwiched between the side chains of Trp56 and Trp102 via cation−*π, π* stacking interactions that positioned the base to form multiple hydrogen bonding interactions with the protein. *N1* and *N2* formed bidentate hydrogen bonds with the carboxylate side chain of Glu103 and carbonyl moiety formed hydrogen bonding interactions with the backbone NH group from Trp102. The phenyl group formed edge-on *π* − *π* stacking with Trp102 placing the phosphonate in a position to form salt-bridge interactions with the amino and guanidine side chains of Lys162 and Arg112, respectively. The *meta*-substituted ethyl linker formed hydrophobic interactions with the side chains of Thr203 and Lys206. The methylene carbon of the *N*-7-methylbenzofuran formed van der Waals interactions with Trp166 and Trp102. The benzofuran of **5-PA** was situated in a hydrophobic pocket lined by Trp166, Val153, Phe48, Leu60 and Pro100 forming edge-on *π* − *π* stacking with Trp56. The chlorine of the benzofuran extended further into a pocket interacting with the hydroxyl side chain Ser92 and the backbone atoms of Phe47 and Phe48. Illumination of the binding mode of **5-PA** occupying the cap-binding domain of eIF4E will be invaluable to the design approach of alternative nucleotide- and non-nucleotide-based derivatives that retain the edge-on-π interaction with W102 while accessing the cationic residues located in the positively charged cleft. Progress with respect to this goal will be reported in due time.

**Figure 5.**
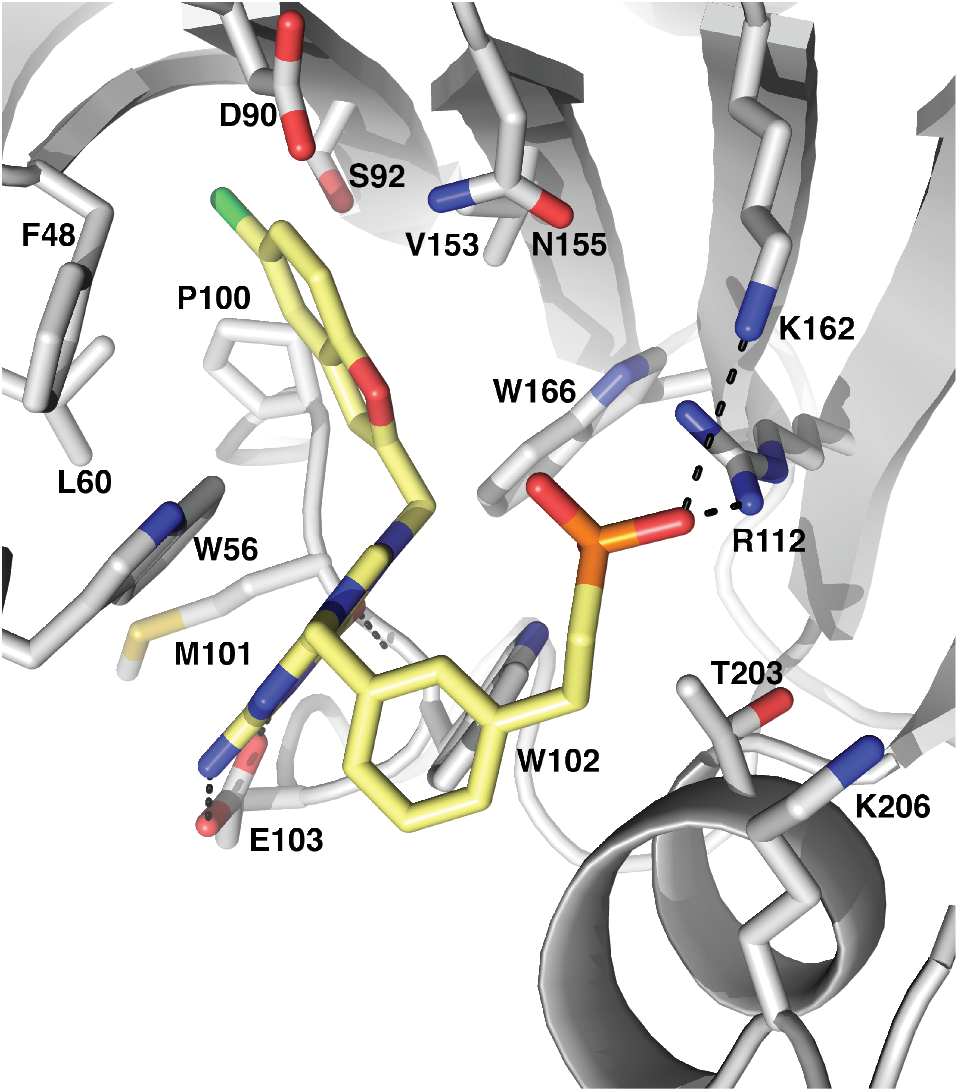
eIF4E bound to **5-PA**. The eIF4E protein backbone is shown as a grey ribbon with residues that interact with **5-PA** shown as sticks with carbons in light yellow. Nitrogen atoms are colored in blue, oxygen in red, chlorine in green, sulfur in gold and phosphorus in orange. Dashed lines represent both hydrogen and ionic bonds. (PDB: 9DON)

To demonstrate that the observed target engagement and anti-proliferative activity in cells was due to on-target inhibition of cap-dependent translation, we examined the impact of **5** on the expression of oncoproteins whose translation is governed by eIF4E. We selected several well-studied mRNAs known to be dependent on eIF4E, including c-Myc, ornithine decarboxylase 1 (ODC1), cyclin D1 (CCND1), and cyclin D3 (CCND3), and used Western blot to visualize their protein levels after treatment of A2058 cells for 6 h. Excitingly, significant decreases were observed in ODC1, CCND1, and CCND3 levels after treatment (Figure 6A). In contrast, there were no significant changes in c-Myc expression levels. This was not surprising, given that Myc translation is driven by both cap-dependent and cap-independent mechanisms due to the presence of an internal ribosome entry site (IRES) in its 5′ UTR.

**Figure 6.**
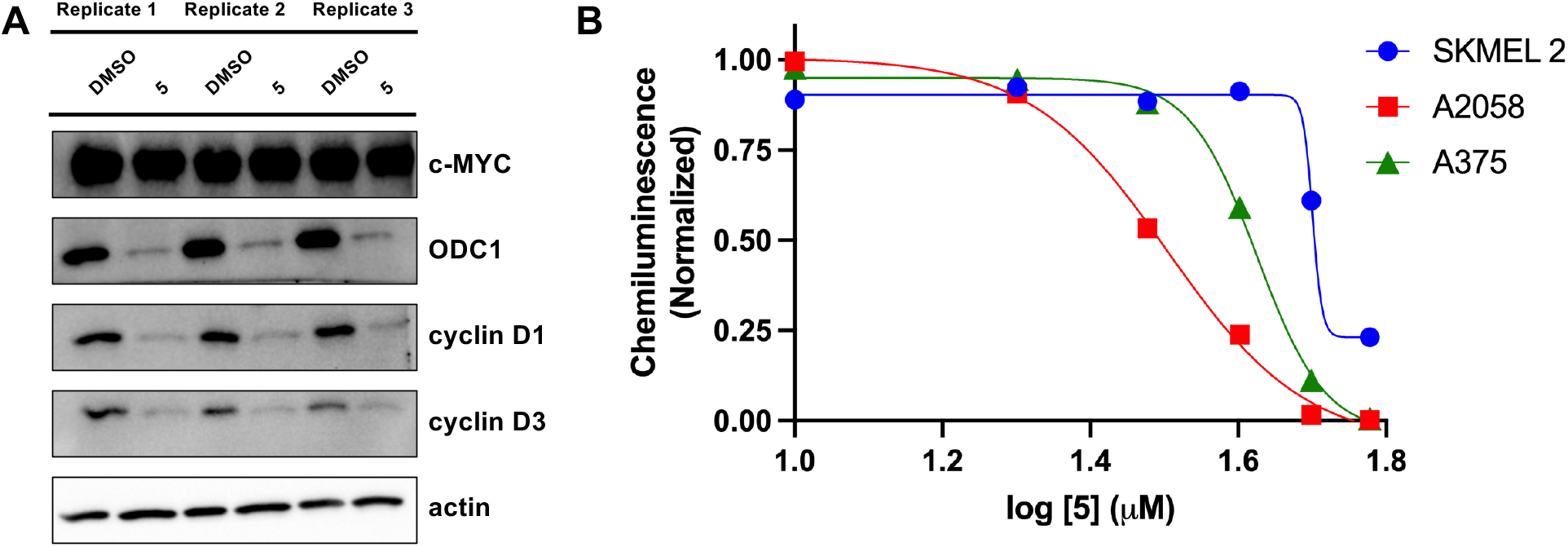
Biological activity of **5** in melanoma cell lines. (A) Effect of **5** on protein levels of select cap-dependent transcripts. A2058 cells were treated for 6 h, and protein levels were measured using Western blot. (B) Dose-dependent anti-proliferative activity of **5** across 3 melanoma cell lines after treatment for 48 h.

Since drug-resistant BRAF^V600E^ melanoma cells have been reported to have a particularly high dependence on eIF4E and cap-dependent translation initiation,^9^ we were interested to compare the anti-proliferative activity of **5** in vemurafenib-resistant and -sensitive melanoma cell lines. As cell lines, we used the drug-resistant BRAF^V600E^ melanoma cell line A2058, the drug-sensitive BRAF^V600E^ cell line A375, and the BRAF-WT cell line SKMEL2. Cells were treated with **5** for 48 h and viability was analyzed using the CellTiter-Glo viability assay. Strikingly, A2058 cells showed the greatest sensitivity to **5**, while the viability of SKMEL2 and A375 cells was only moderately affected (Figure 6B). Importantly, these findings phenocopy the results of this previous report and demonstrate that multiple pharmacological mechanisms of eIF4F inhibition, including targeting the eIF4E/eIF4G PPI, eIF4A helicase activity, and now eIF4E−m^7^GpppX capped mRNA binding, can be utilized for inhibiting cap-dependent translation and overcoming drug resistance in vemurafenib-resistant melanoma.^9^ In sum, these results provide additional validation for eIF4E as a promising target for re-sensitizing drug-resistant melanoma cells to existing treatments and the potential of cap analogues or other eIF4E cap binding site-targeted agents for therapeutically addressing this hypothesis.

## Conclusion

The development of resistance to current melanoma therapies has been a major hindrance to successfully treating this deadly form of skin cancer. Drug-resistant melanoma has been shown to be highly dependent on the overexpression of eIF4E and enhanced cap-dependent translation. Thus, eIF4E has arisen as an attractive therapeutic target for re-sensitizing drug-resistant melanoma to existing therapies. Expanding on our previous work of designing cell-permeable cap analogue prodrugs that inhibit eIF4E binding to capped mRNA, we designed a series of second-generation cap analogues exploring the effects of alternative *N*-9-substituted linkers between the nucleobase and phosphonate moieties on biochemical and cellular potency. *Meta*-substituted *N*-9-benzyl analogue **5**, the most potent in the series, displayed similar eIF4E inhibitory activity and cellular antiproliferative activity to our first-generation compounds. Importantly, drug-resistant melanoma cells showed greater sensitivity to cap analogue **5** than drug-sensitive cells, indicating that melanoma treatment may benefit from the inclusion of eIF4E inhibitors to overcome drug resistance. Ongoing efforts are focused on the development of new analogues with optimized potentcy and *in vivo* drug-like properties to address this question and will be reported in due course.

## ASSOCIATED CONTENT

### Supporting Information

The Supporting Information is available free of charge at…

Methods, supplemental schemes and figures, NMR spectra, HPLC spectra, and crystallography data collection and refinement statistics (PDF)

Molecular formula strings (PDF)

### Accession Codes

eIF4E bound to **5-PA**, PDB: 9DON

## AUTHOR INFORMATION

### Authors

**Emilio L. Cárdenas** - Department of Medicinal Chemistry, College of Pharmacy, University of Michigan, 1600 Huron Parkway, NCRC B520, Ann Arbor, Michigan 48109, USA

**Rachel L. O’Rourke** - Department of Medicinal Chemistry, College of Pharmacy, University of Michigan, 1600 Huron Parkway, NCRC B520, Ann Arbor, Michigan 48109, USA

**Arya Menon** - Department of Medicinal Chemistry, College of Pharmacy, University of Michigan, 1600 Huron Parkway, NCRC B520, Ann Arbor, Michigan 48109, USA

**Gabriela Vega-Hernández** - Program in Chemical Biology, University of Michigan, Ann Arbor, Michigan 48109, United States

**Jennifer Meagher -** Life Science Institute, University of Michigan, Ann Arbor, Michigan 48109, United States.

**Jeanne Stuckey** - Life Science Institute, University of Michigan, Ann Arbor, Michigan 48109, United States.

## ACKNOWLEDGMENTS

This work was supported by the NIH (R01 GM132341 to A.L.G.) and the Rogel Cancer Center. This research used resources of the Advanced Photon Source, a U.S. Department of Energy (DOE) Office of Science User Facility operated for the DOE Office of Science by Argonne National Laboratory under Contract No. DE-AC02-06CH11357. Use of the LS-CAT Sector 21 was supported by the Michigan Economic Development Corporation and the Michigan Technology Tri-Corridor (Grant 085P1000817). This study is supported, in part, by the Cancer Center Support Grant (P30CA046592) from the National Cancer Institute, National Institutes of Health, to Rogel Cancer Center of the University of Michigan.

## ABBREVIATIONS

eIF4E,: eukaryotic translation initiation factor 4E;
POM,: pivaloyloxymethyl;
FP,: fluorescence polarization;
ODC1,: ornithine decarboxylase 1;
UTR,: untranslated region.

## TABLE OF CONTENTS GRAPHIC

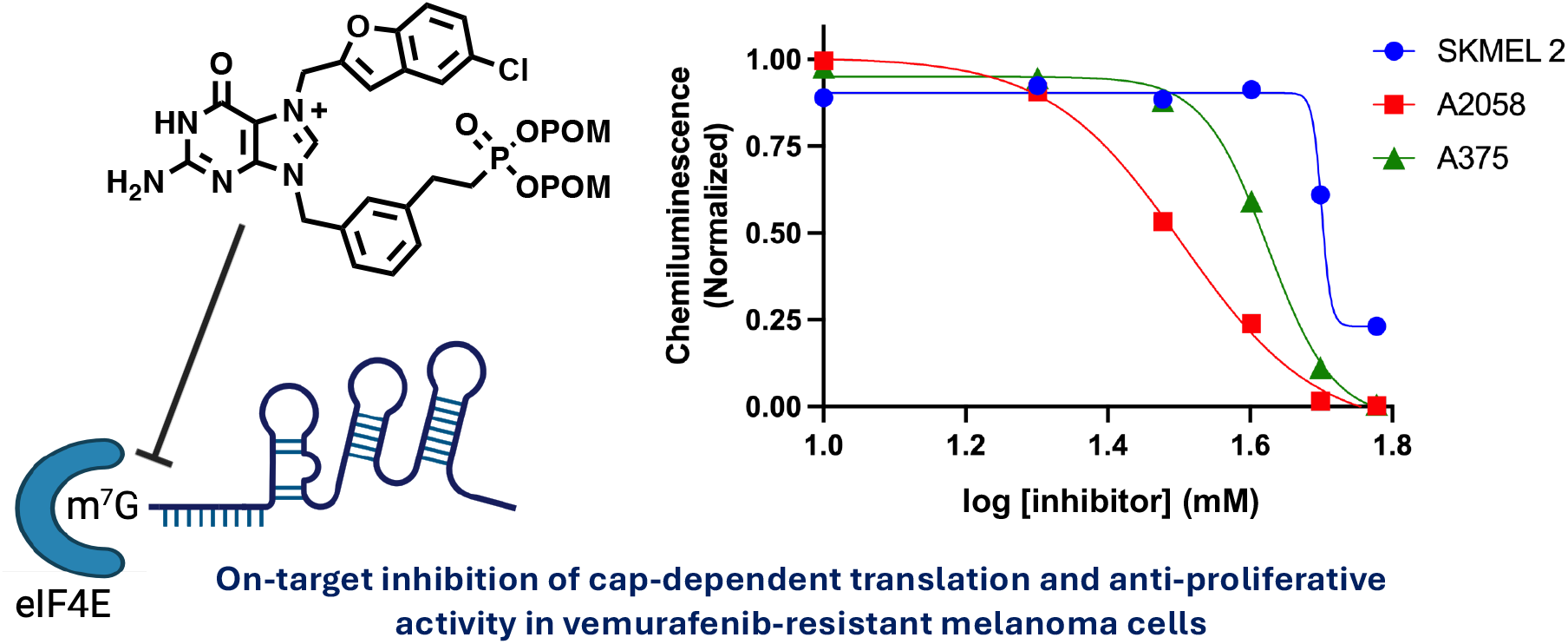

## REFERENCES

1. Rebecca, V. W.; Herlyn, M., Nongenetic mechanisms of drug resistance in melanoma. Annu. Rev. Cancer Biol. 2020, 4, 315–330.

2. Winder, M.; Viros, A., Mechanisms of drug resistance in melanoma. In Mechanisms of drug resistance in cancer therapy, Mandala, M.; Romano, E., Eds. Springer: 2017; Vol. 249, pp 91–108.

3. Patton, E. E.; Mueller, K. L.; Adams, D. J.; Anandasabapthy, N.; Aplin, A. E.; Bertolotto, C.; Bosenberg, M.; Ceol, C. J.; Burd, C. E.; Chi, P.; Herlyn, M.; Holmen, S. L.; Karreth, F. A.; Kaufman, C. K.; Khan, S.; Kobold, S.; Leucci, E.; Levy, C.; Lombard, D. B.; Lund, A. W.; Marie, K. L.; Marine, J.-L.; Marais, R.; McMahon, M.; Robles-Espinoza, C. D.; Ronai, Z. A.; Samuels, Y.; Soengas, M. S.; Villanueva, J.; Weeraratna, A. T.; White, R. M.; Yeh, I.; Zhu, J.; Zon, L. I.; Hurlbert, M. S.; Merlino, G., Melanoma models for the next generation of therapies. Cancer Cell 2021, 39, 610–631.

4. Kalaora, S.; Nagler, A.; Wargo, J. A.; Samuels, Y., Mechanisms of immune activation and regulation: lessons from melanoma. Nat. Rev. Cancer 2022, 22, 195–207.

5. Fiskus, W.; Mitsiades, N., B-Raf inhibition in the clinic: present and future. Annu. Rev. Med. 2016, 67, 29–43.

6. Khosravi, S.; Tam, K. J.; Ardekani, G. S.; Martinka, M.; McElwee, K. J.; Ong, C. J., eIF4E is an adverse prognostic marker of melanoma patient survival by increasing melanoma cell invasion. J. Invest. Dermatol. 2015, 135, 1358–1367.

7. Carter, J. H.; Deddens, J. A.; Spaulding IV, N. R.; Lucas, D.; Colligan, B. M.; Lewis, T. G.; Hawkins, E.; Jones, J.; Pemberton, J. O.; Douglass, L. E.; Graff, J. R., Phosphorylation of eIF4E serine 209 is associated with tumour progression and reduced survival in malignant melanoma. Br. J. Cancer 2016, 114, 444–453.

8. O’Reilly, K. E.; Warycha, M.; Davies, M. A.; Rodrik, V.; Zhou, X. K.; Yee, H.; Polsky, D.; Pavlick, A. C.; Rosen, N.; Bhardwaj, N. B.; Mills, G.; Osman, I., Phosphorylated 4E-BP1 is associated with poor survival in melanoma. Clin. Cancer Res. 2009, 15, 2872–2878.

9. Boussemart, L.; Malka-Mahieu, H.; Girault, I.; Allard, D.; Hemmingsson, O.; Tomasic, G.; Thomas, M.; Basmadjian, C.; Ribeiro, N.; Thuaud, F.; Mateus, C.; Routier, E.; Kamsu-Kom, N.; Agoussi, S.; Eggermont, A. M.; Desaubry, L.; Robert, C.; Vagner, S., eIF4F is a nexus of resistance to anti-BRAF and anti-MEK cancer therapies. Nature 2014, 513, 105–109.

10. Zhan, Y.; Dahabieh, M. S.; Rajakumar, A.; Dobocan, M. C.; M’Boutchou, M.-N.; Goncalves, C.; Shiru, L.; Pettersson, F.; Topisirovic, I.; van Kempen, L.; del Rincon, S. V.; Miller Jr., W. H., The role of eIF4E in response and acquired resistance to vemurafenib in melanoma. J. Invest. Dermatol. 2015, 135, 1368–1376.

11. Cerezo, M.; Guemiri, R.; Druillennec, S.; Girault, I.; Malka-Mahieu, H.; Shen, S.; Allard, D.; Martineau, S.; Welsch, C.; Agoussi, S.; Estrada, C.; Adam, J.; Libenciuc, C.; Routier, E.; Roy, S.; Desaubry, L.; Eggermont, A. M.; Sonenberg, N.; Scoazec, J. Y.; Eychene, A.; Vagner, S.; Robert, C., Translational control of tumor immune escape via the eIF4F-STAT1-PD-L1 axis in melanoma. Nat. Med. 2018, 24, 1877–1886.

12. Huang, F.; Goncalves, C.; Bartish, M.; Remy-Sarrazin, J.; Issa, M. E.; Cordeiro, B.; Guo, Q.; Emond, A.; Attias, M.; Yang, W.; Plourde, D.; Su, J.; Gimeno, M. G.; Zhan, Y.; Galan, A.; Rzymski, T.; Mazan, M.; Masiejczyk, M.; Faber, J.; Khoury, E.; Benoit, A.; Gagnon, N.; Dankort, D.; Journe, F.; Ghanem, G. E.; Krawczyk, C. M.; Saragovi, H. U.; Piccirillo, C. A.; Sonenberg, N.; Topisirovic, I.; Rudd, C. E.; Miller, W. H., Jr.; Del Rincon, S. V., Inhibiting the MNK1/2-eIF4E axis impairs melanoma phenotype switching and potentiates antitumor immune responses. J. Clin. Invest. 2021, 131, e140752.

13. Cardenas, E. L.; O’Rourke, R. L.; Menon, A.; Meagher, J.; Stuckey, J.; Garner, A. L., Design of cell-permeable inhibitirs of eukaryotic translation initiation factor 4E (eIF4E) for inhibiting aberrant cap-dependent translation in cancer. J. Med. Chem. 2023, 66, 10734–10745.

14. Cardenas, E. L.; O’Rourke, R. L.; Menon, A.; Meagher, J.; Stuckey, J.; Garner, A. L., Design of Cell-Permeable Inhibitors of Eukaryotic Translation Initiation Factor 4E (eIF4E) for Inhibiting Aberrant Cap-Dependent Translation in Cancer. J Med Chem 2023, 66 (15), 10734–10745.

15. Chen, X.; Kopecky, D. J.; Mihalic, J.; Jeffries, S.; Min, X.; Heath, J.; Deignan, J.; Lai, S.; Fu, Z.; Guimaraes, C.; Shen, S.; Li, S.; Johnstone, S.; Thibault, S.; Xu, H.; Cardozo, M.; Shen, W.; Walker, N.; Kayser, F.; Wang, Z., Structure-guided design, synthesis, and evaluation of guanine-derived inhibitors of the eIF4E mRNA-cap interaction. J. Med. Chem. 2012, 55, 3837–3851.

16. Halazy, S.; Eggenspiller, A.; Ehrhard, A.; Danzin, C., Phosphonate derivatives of N9-benzylguanine: a new class of potent purine nucleoside phosphorylase inhibitors. Bioorg. Med. Chem. Lett. 1992, 2, 407–410.

17. Pautus, S.; Sehr, P.; Lewis, J.; Fortune, A.; Wolkerstorfer, A.; Szolar, O.; Guilligay, D.; Lunardi, T.; Decout, J.-L.; Cusack, S., New 7-Methylguanine derivatives targeting the influenza polymerase PB2 cap-binding domain. J. Med. Chem. 2013, 56, 8915–8930.

18. Visco, C.; Perrera, C.; Thieffine, S.; Sirtori, F. R.; D’Allesio, R.; Magnaghi, P., Development of biochemical assays for the identification of eIF4E-specific inhibitors. J. Biomol. Screen. 2012, 17, 581–592.

19. Molina, D. M.; Jafari, R.; Ignatushchenko, M.; Seki, T.; Larsson, E. A.; Dan, C.; Sreekumar, L.; Cao, Y.; Nordlund, P., Monitoring drug target engagement in cells and tissues using the cellular thermal shift assay. Science 2013, 341, 84–87.

