## Supporting Information for "Second-Generation Cap Analogue Prodrugs for Targeting Aberrant Eukaryotic Translation Initiation Factor 4E (eIF4E) Activity in Drug-Resistant Melanoma"

[1]Department of Medicinal Chemistry, College of Pharmacy, University of Michigan, 1600 Huron Parkway, NCRC B520, Ann Arbor, Michigan 48109

**A. Characterization Data for Compounds 2−7** Pages S2−S15

**B. General Methods** Pages S16−S21

**C. Synthetic Methods** Pages S21−S32

**D. X-ray Crystallography Data** Page S33−S34

**E. References** Page S35

#### A. Characterization Data for Compounds 2–7

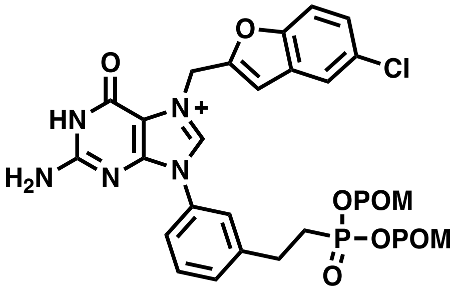


*2-amino-9-(3-(2-(bis((pivaloyloxy)methoxy)phosphoryl)ethyl)phenyl)-7-((5-chlorobenzofuran-2-yl)methyl)-6-oxo-6,9-dihydro-1H-purin-7-ium (**2**)*. $^{1}$H NMR (500 MHz, DMSO-$d_6$) δ 9.79 (s, 1H), 7.77 (d, $J$ = 2.3 Hz, 1H), 7.64 (t, $J$ = 5.4 Hz, 2H), 7.61 (s, 1H), 7.57 (t, $J$ = 8.0 Hz, 1H), 7.48 (d, $J$ = 7.8 Hz, 1H), 7.37 (d, $J$ = 8.8, 2.2 Hz, 1H), 7.15 (s, 1H), 5.91 (s, 2H), 5.61 (m, 4H), 2.88 (m, 2H), 2.26 (m, 2H), 1.15 (s, 18H); $^{13}$C NMR (125 MHz, DMSO-$d_6$) δ 176.63, 153.50, 152.26, 150.64, 142.62, 142.47, 137.53, 133.02, 130.12, 130.08, 129.77, 128.12, 125.51, 125.22, 123.87, 121.59, 113.43, 107.59, 107.09, 81.88, 81.84, 45.36, 38.66, 27.72, 27.63, 26.93, 26.54; LRMS (ESI$^{+}$) 728.2769 [M]$^{+}$.

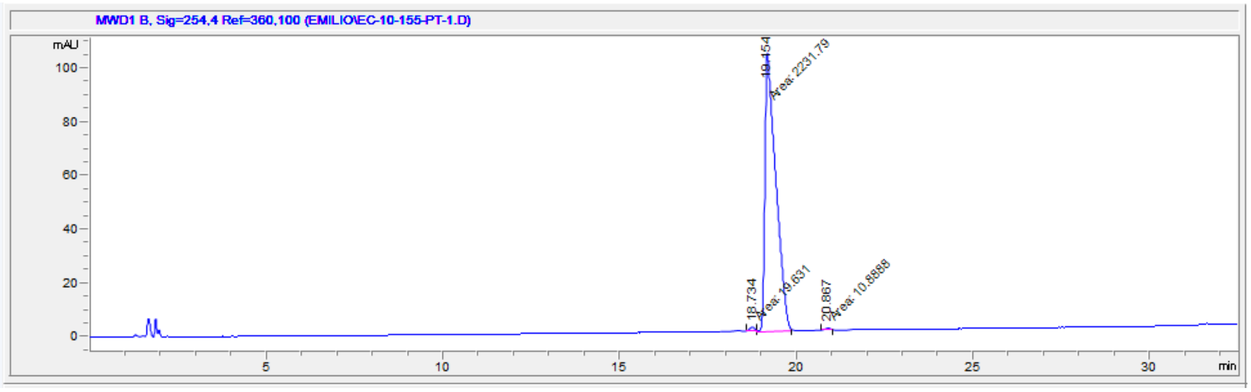


| # | Time | Area | Height | Width | Area% | Symmetry |
|---|---|---|---|---|---|---|
| 1 | 18.734 | 19.6 | 1.8 | 0.1841 | 0.868 | 1.085 |
| 2 | 19.154 | 2231.8 | 104.3 | 0.3567 | 98.651 | 0.27 |
| 3 | 20.867 | 10.9 | 9E-1 | 0.2019 | 0.481 | 1.032 |

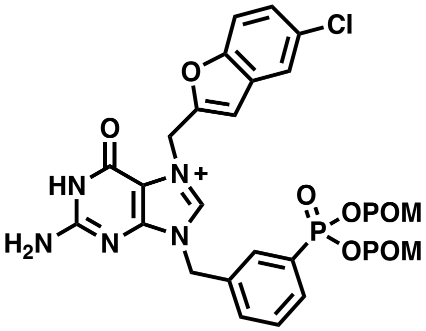


*2-amino-9-(3-(bis((pivaloyloxy)methoxy)phosphoryl)benzyl)-7-((5-chlorobenzofuran-2-yl)methyl)-6-oxo-6,9-dihydro-1H-purin-7-ium (**3**)*. $^{1}$H NMR (500 MHz, DMSO-d$_6$) δ 9.14 (s, 1H), 8.41 (bs, 1H), 7.82 (d, *J* = 14.5 Hz, 1H), 7.73 (d, *J* = 2.2 Hz, 1H), 7.68 (d, *J* = 7.8 Hz, 1H), 7.63 (m, 1H), 7.57 (d, *J* = 8.7 Hz, 1H), 7.56 (m, 1H), 7.33 (dd, *J* = 8.7, 2.1 Hz, 1H), 7.03 (s, 1H), 5.90 (s, 2H), 5.64 (m, 4H), 0.95 (s, 18H); $^{13}$C NMR (125 MHz, DMSO-d$_6$) δ 176.34, 153.35, 136.42, 133.13, 129.80, 128.04, 125.32, 121.49, 113.28, 107.07, 82.21, 40.90, 38.48, 26.66; LRMS (ESI$^{+}$) 714.2401 [M]$^{+}$.

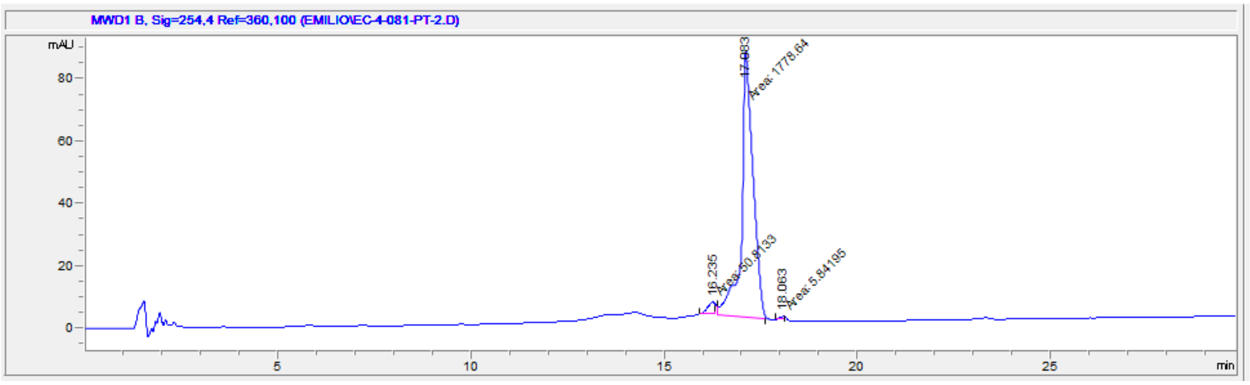


| # | Time | Area | Height | Width | Area% | Symmetry |
|---|---|---|---|---|---|---|
| 1 | 16.235 | 50.8 | 3.6 | 0.2323 | 2.769 | 1.634 |
| 2 | 17.083 | 1778.6 | 85.1 | 0.3482 | 96.913 | 0.608 |
| 3 | 18.063 | 5.8 | 8.2E-1 | 0.1188 | 0.318 | 1.908 |

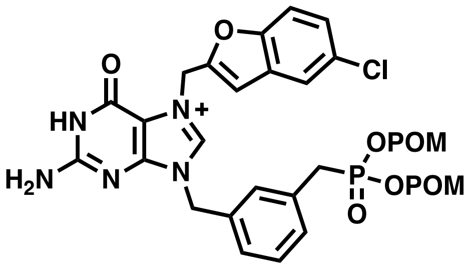


*2-amino-9-(3-((bis((pivaloyloxy)methoxy)phosphoryl)methyl)benzyl)-7-((5-chlorobenzofuran-2-yl)methyl)-6-oxo-6,9-dihydro-1H-purin-7-ium (**4**)*. $^{1}$H NMR (400 MHz, MeOD-$d_4$) δ 7.61 (d, *J* = 2.7 Hz, 1H), 7.45 (m, 2H), 7.44 (s, 1H), 7.38 (t, *J* = 9.8 Hz, 1H), 7.31 (m, 1H), 7.30 (dd, *J* = 11.0, 2.7 Hz, 1H), 7.09 (s, 1H), 5.88 (s, 2H), 5.57 (s, 4H), 5.42 (s, 2H), 3.37 (d, *J* = 27.7 Hz, 2H), 1.15 (s, 18H); $^{13}$C NMR (125 MHz, MeOD-$d_4$) 176.70, 153.69, 151.25, 130.32, 130.26, 129.87, 129.31, 129.17, 128.62, 127.36, 125.19, 120.89, 112.19, 107.13, 81.63, 81.58, 44.67, 38.24, 33.35, 32.25, 25.73; LRMS (ESI$^{+}$) 728.2715 [M]$^{+}$.

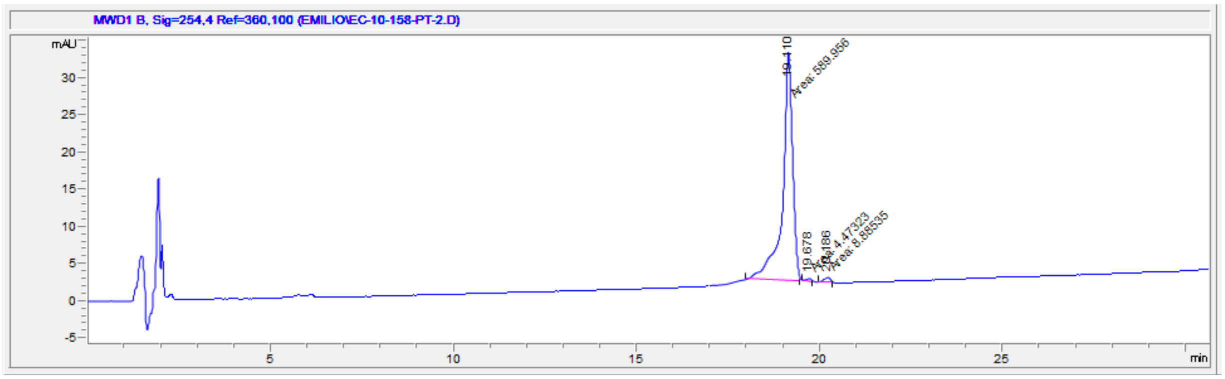


| # | Time | Area | Height | Width | Area% | Symmetry |
|---|---|---|---|---|---|---|
| 1 | 19.11 | 590 | 30.8 | 0.3193 | 97.786 | 1.369 |
| 2 | 19.678 | 4.5 | 4.2E-1 | 0.1759 | 0.741 | 0.775 |
| 3 | 20.186 | 8.9 | 6.9E-1 | 0.2152 | 1.473 | 1.476 |

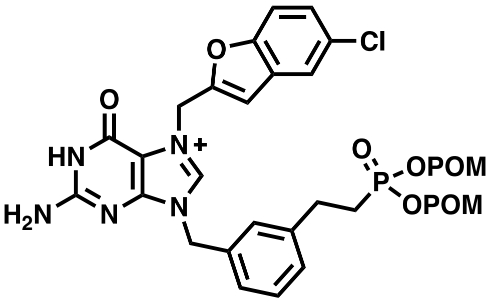


*2-amino-9-(3-(2-(bis((pivaloyloxy)methoxy)phosphoryl)ethyl)benzyl)-7-((5-chlorobenzofuran-2-yl)methyl)-6-oxo-6,9-dihydro-1H-purin-7-ium (**5**)*. $^{1}$H NMR (500 MHz, DMSO-$d_6$) δ 9.42 (s, 1H), 7.76 (d, *J* = 2.2 Hz, 1H), 7.59 (d, *J* = 8.7 Hz, 1H), 7.35 (dd, *J* = 8.9, 2.2 Hz, 1H), 7.32 (d, *J* = 7.6 Hz, 1H), 7.24 (dd, *J* = 15.6, 7.8 Hz, 2H), 7.07 (s, 1H), 5.87 (s, 2H), 5.59 (m, 4H), 5.33 (s, 2H), 2.77 (m, 1H), 2.18 (m, 1H), 1.13 (s, 18H); $^{13}$C NMR (125 MHz, DMSO-$d_6$) δ 176.61, 153.45, 152.50, 150.44, 141.50, 141.35, 137.70, 135.07, 129.71, 129.51, 128.51, 128.14, 128.02, 126.47, 125.52, 121.60, 113.33, 107.40, 107.20, 81.84, 81.79, 48.01, 45.17, 38.64, 27.80, 27.77, 27.73, 26.91, 26.64; LRMS (ESI$^{+}$) 742.2529 [M]$^{+}$.

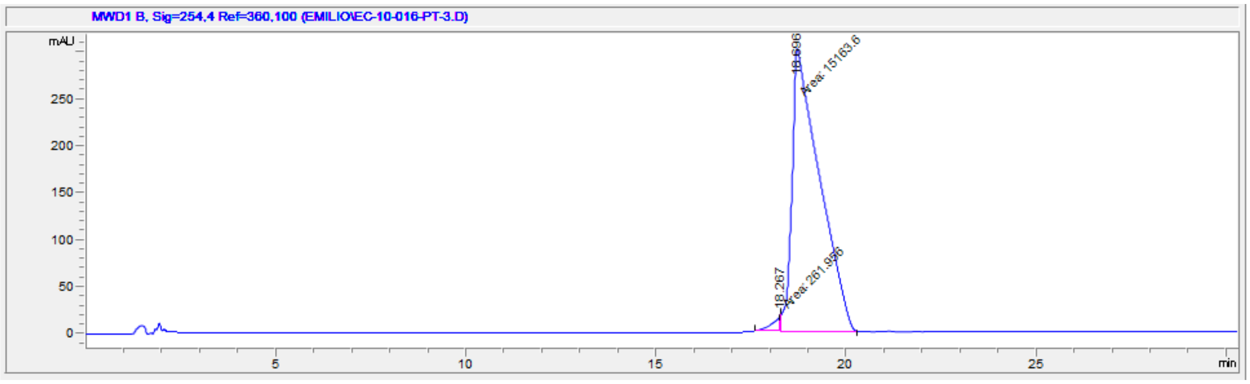


| # | Time | Area | Height | Width | Area% | Symmetry |
|---|---|---|---|---|---|---|
| 1 | 18.267 | 262 | 16.8 | 0.2601 | 1.698 | 0 |
| 2 | 18.696 | 15163.6 | 301.5 | 0.8381 | 98.302 | 0.236 |

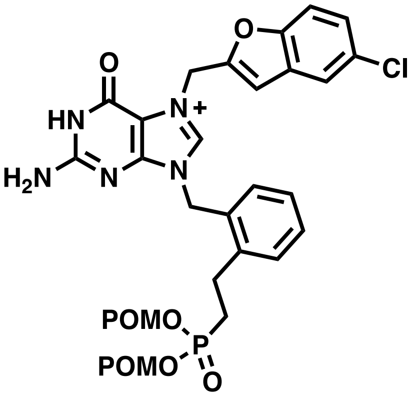


*2-amino-9-(2-(2-(bis((pivaloyloxy)methoxy)phosphoryl)ethyl)benzyl)-7-((5-chlorobenzofuran-2-yl)methyl)-6-oxo-6,9-dihydro-1H-purin-7-ium (**6**)*. $^1$H NMR (500 MHz, DMSO-$d_6$) δ 9.41 (s, 1H), 7.76 (d, $J$ = 2.2 Hz, 1H), 7.59 (d, $J$ = 8.6 Hz, 1H), 7.35 (dd, $J$ = 8.9, 2.4 Hz, 1H), 7.33 (d, $J$ = 4.0 Hz, 2H), 7.25 (m, 1H), 7.10 (d, $J$ = 7.6 Hz, 1H), 7.09 (s, 1H), 5.87 (s, 2H), 5.60 (m, 4H), 5.41 (s, 2H), 2.95 (m, 2H), 2.24 (m, 2H), 1.13 (s, 18H); LRMS (ESI$^+$) 742.2391 [M]$^+$.

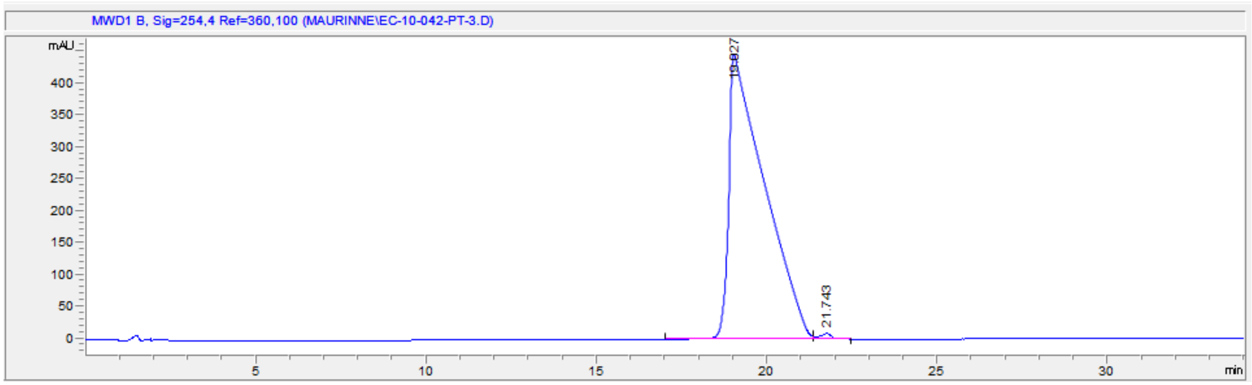


| # | Time | Area | Height | Width | Area% | Symmetry |
|---|---|---|---|---|---|---|
| 1 | 19.027 | 32711.4 | 448.6 | 0.9168 | 99.364 | 0.17 |
| 2 | 21.743 | 209.4 | 8.7 | 0.3521 | 0.636 | 1.454 |

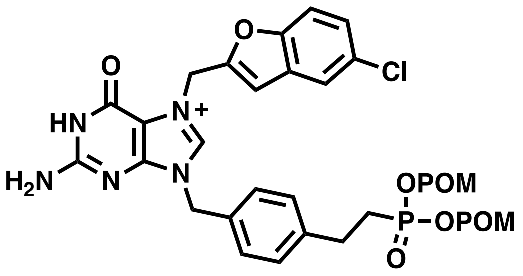


*2-amino-9-(4-(2-(bis((pivaloyloxy)methoxy)phosphoryl)ethyl)benzyl)-7-((5-chlorobenzofuran-2-yl)methyl)-6-oxo-6,9-dihydro-1H-purin-7-ium (7).* $^{1}$H NMR (500 MHz, DMSO-$d_6$) δ 9.55 (s, 1H), 7.76 (d, *J* = 2.2 Hz, 1H), 7.60 (d, *J* = 8.6 Hz, 1H), 7.37 (d, *J* = 2.3 Hz, 1H), 7.34 (d, *J* = 7.9 Hz, 2H), 7.27 (d, *J* = 7.9 Hz, 2H), 7.10 (s, 1H), 5.85 (s, 2H), 5.59 (m, 4H), 5.34 (s, 2H), 2.77 (m, 2H), 2.18 (m, 2H), 1.15 (s, 18H); $^{13}$C NMR (125 MHz, DMSO-$d_6$) δ 176.61, 156.72, 154.23, 153.47, 152.20, 150.27, 141.21, 141.07, 138.49, 132.83, 129.69, 128.97, 128.65, 128.15, 125.57, 121.64, 113.37, 107.60, 107.18, 81.83, 81.79, 48.04, 45.34, 38.65, 27.78, 27.63, 27.59, 26.94, 26.69; LRMS (ESI$^+$) 742.2408 [M]$^+$.

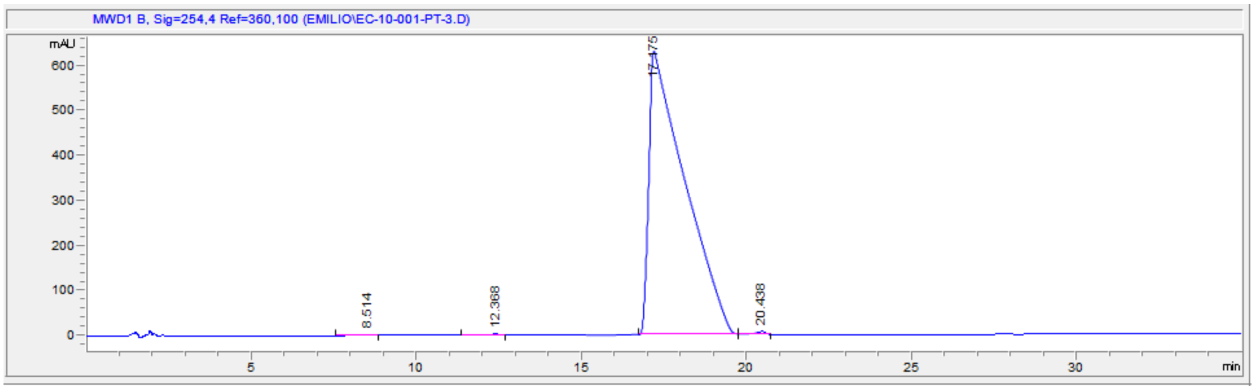


| # | Time | Area | Height | Width | Area% | Symmetry |
|---|---|---|---|---|---|---|
| 1 | 8.514 | 53.6 | 2.6 | 0.2841 | 0.112 | 2.375 |
| 2 | 12.368 | 66.9 | 3.1 | 0.2936 | 0.140 | 2.938 |
| 3 | 17.175 | 47516.2 | 628.8 | 0.9317 | 99.466 | 0.157 |
| 4 | 20.438 | 134.5 | 6.6 | 0.3055 | 0.282 | 1.754 |

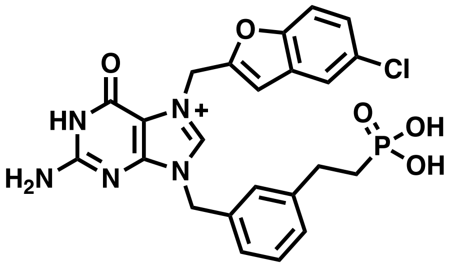


*2-amino-7-((5-chlorobenzofuran-2-yl)methyl)-6-oxo-9-(3-(2-phosphonoethyl)benzyl)-6,9-dihydro-1H-purin-7-ium (**5-PA**)*. LRMS (ESI $^+$) 514.1027 [M]$^+$.

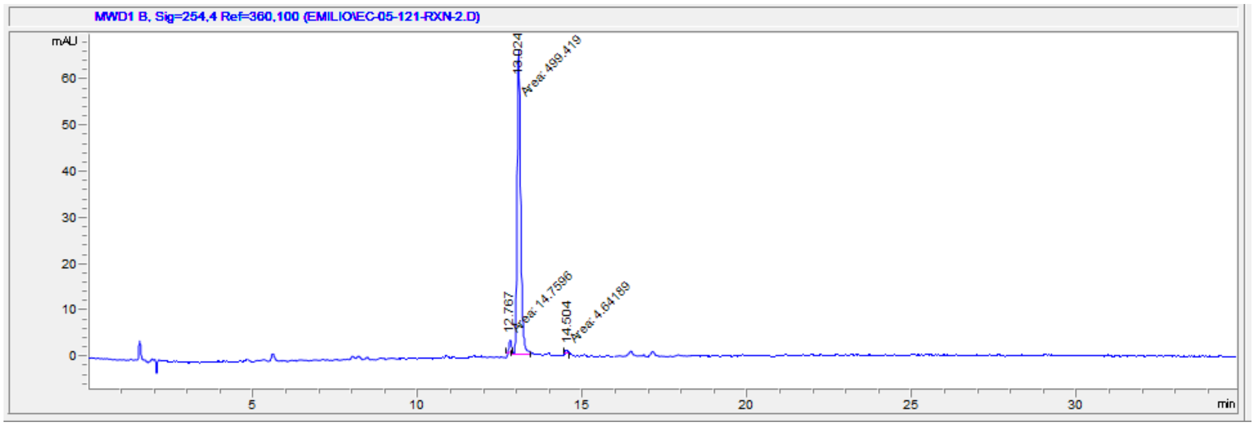


| # | Time | Area | Height | Width | Area% | Symmetry |
|---|---|---|---|---|---|---|
| 1 | 12.767 | 14.8 | 2.9 | 0.0849 | 2.845 | 0.938 |
| 2 | 13.024 | 499.4 | 66 | 0.126 | 96.260 | 0.762 |
| 3 | 14.504 | 4.6 | 8.9E-1 | 0.0871 | 0.895 | 0.927 |

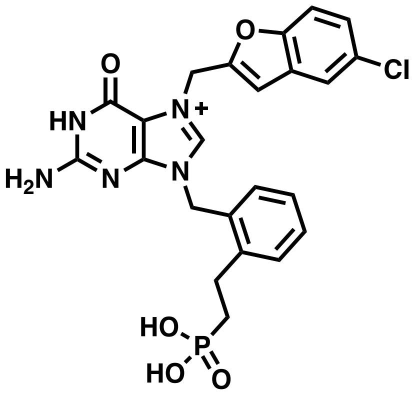


*2-amino-7-((5-chlorobenzofuran-2-yl)methyl)-6-oxo-9-(2-(2-phosphonoethyl)benzyl)-6,9-dihydro-1H-purin-7-ium (**6-PA**)*. LRMS (ESI $^+$) 514.1040 $[M]^+$.

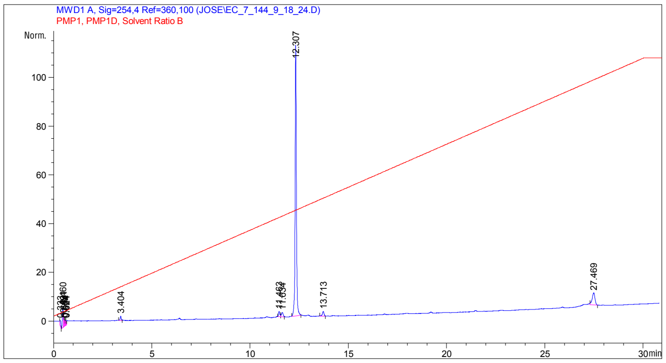


**Table S1.** Analytical purity of inhibitors **2–7**, **5-PA**, **6-PA** determined by RP-HPLC

| **Inhibitor** | **Purity (%)** |
|---|---|
| **2** | 98.7 |
| **3** | 96.9 |
| **4** | 97.8 |
| **5** | 98.3 |
| **6** | 99.3 |
| **7** | 99.4 |
| **5-PA** | 96.3 |
| **6-PA** | 92.8 |

### NMR Spectra for Compounds 2−7

#### Compound 2

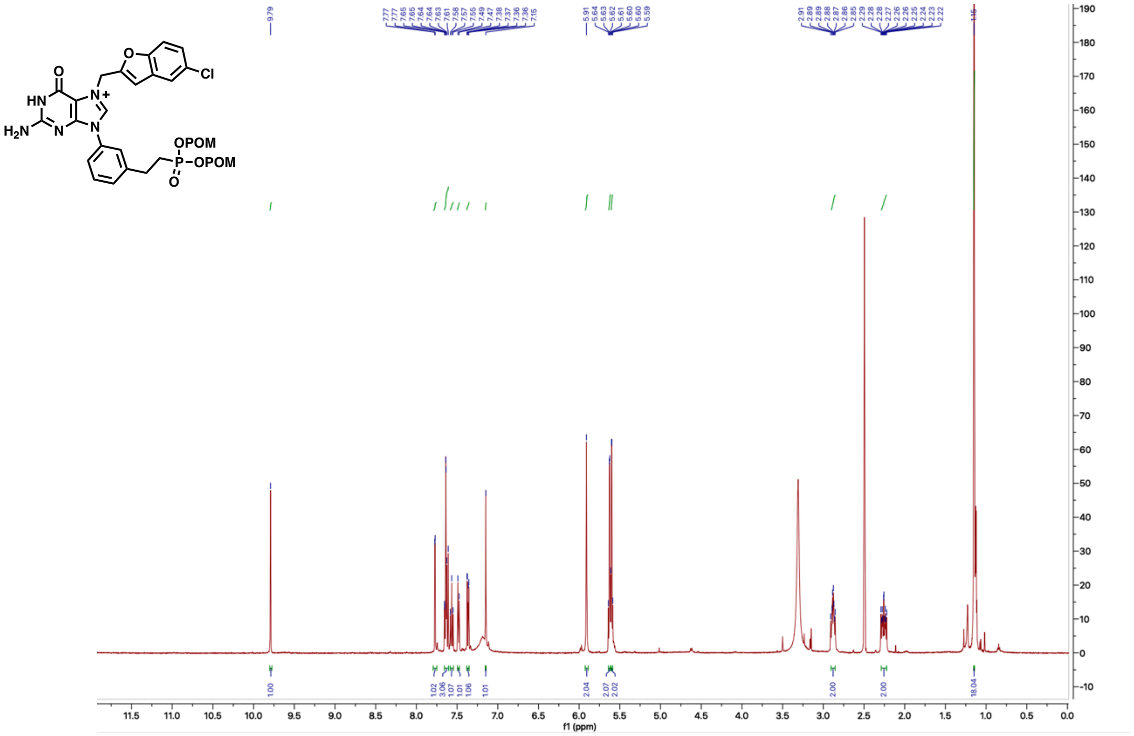


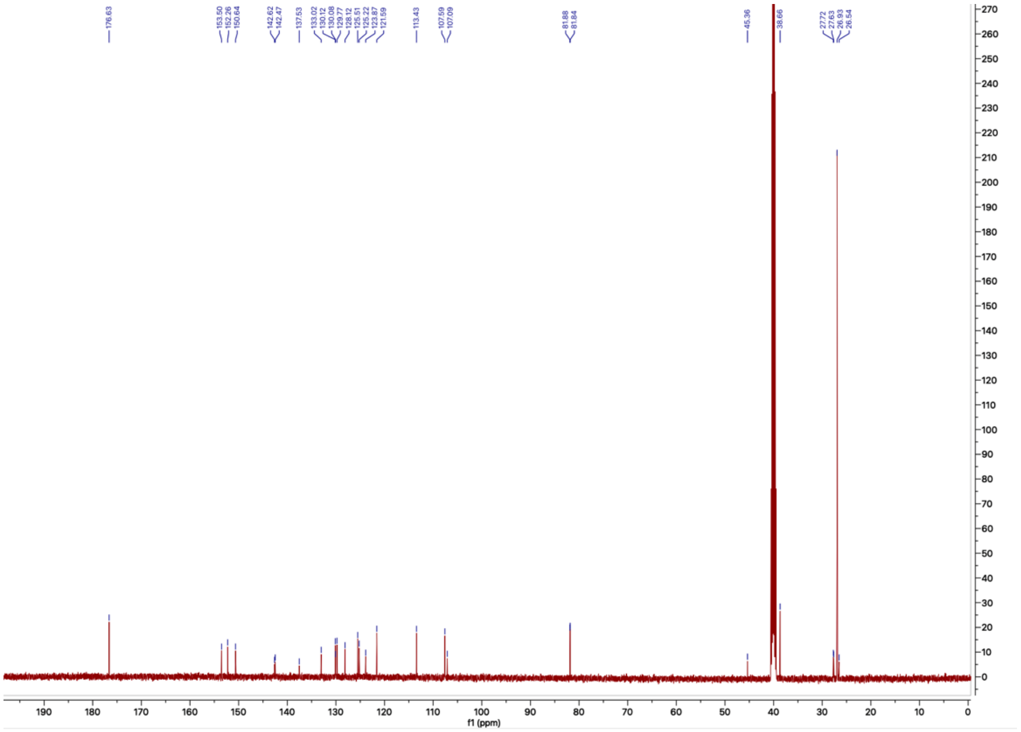

#### Compound 3

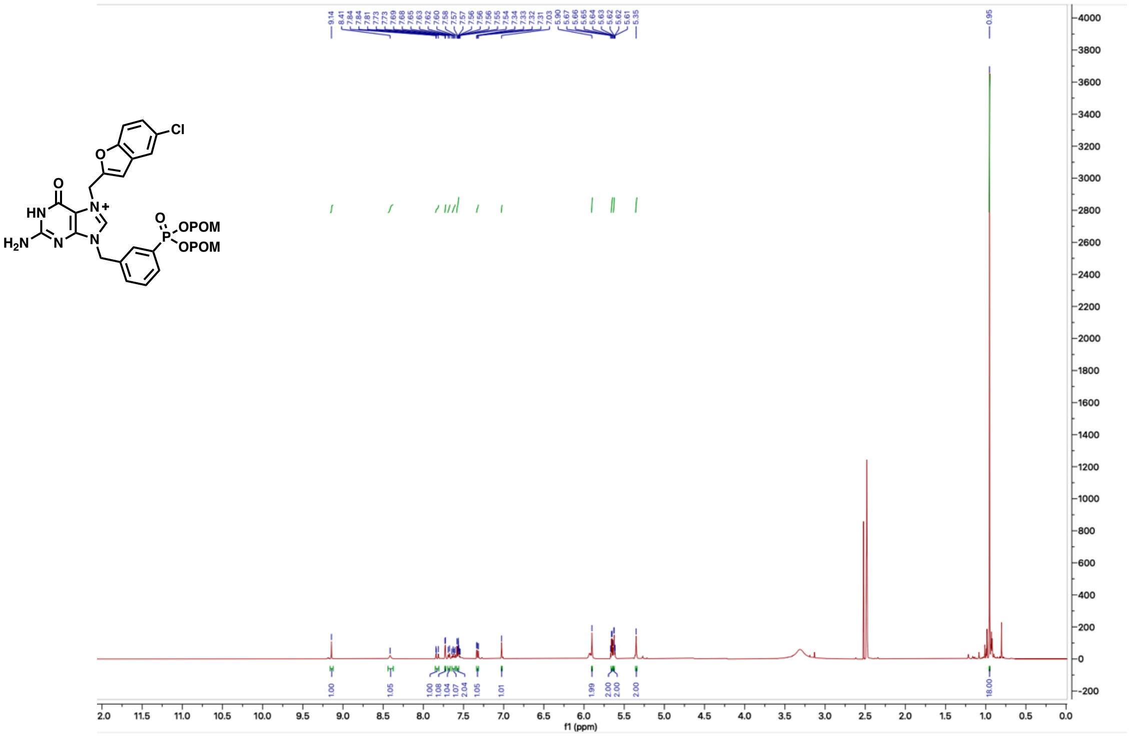

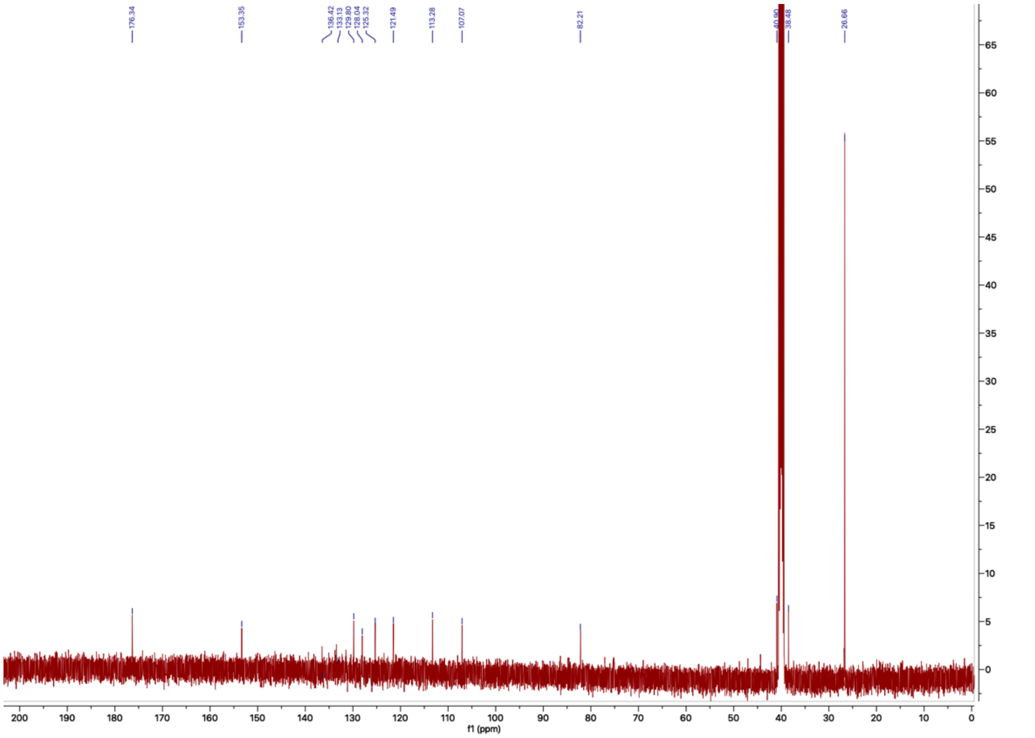

**Compound 4**

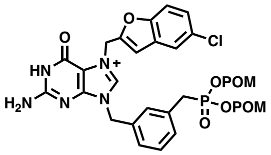


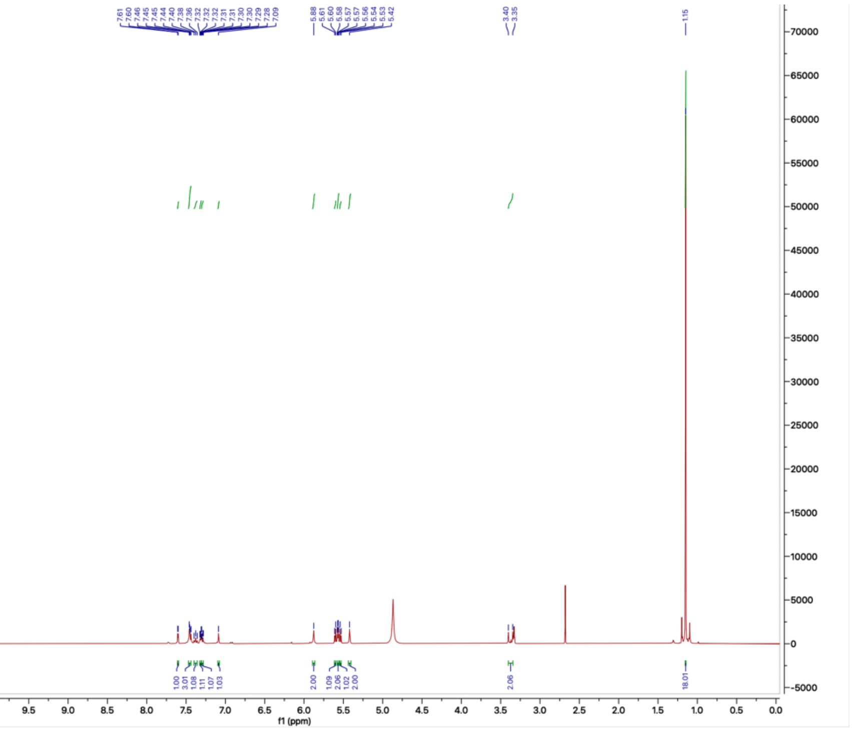

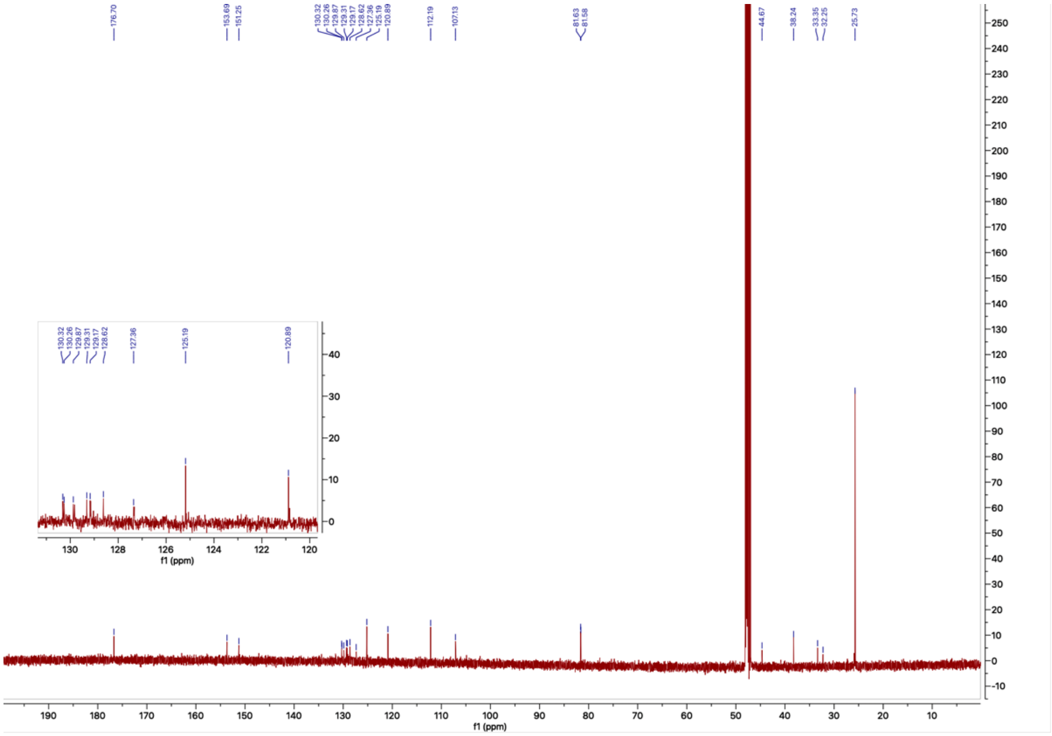

#### Compound 5

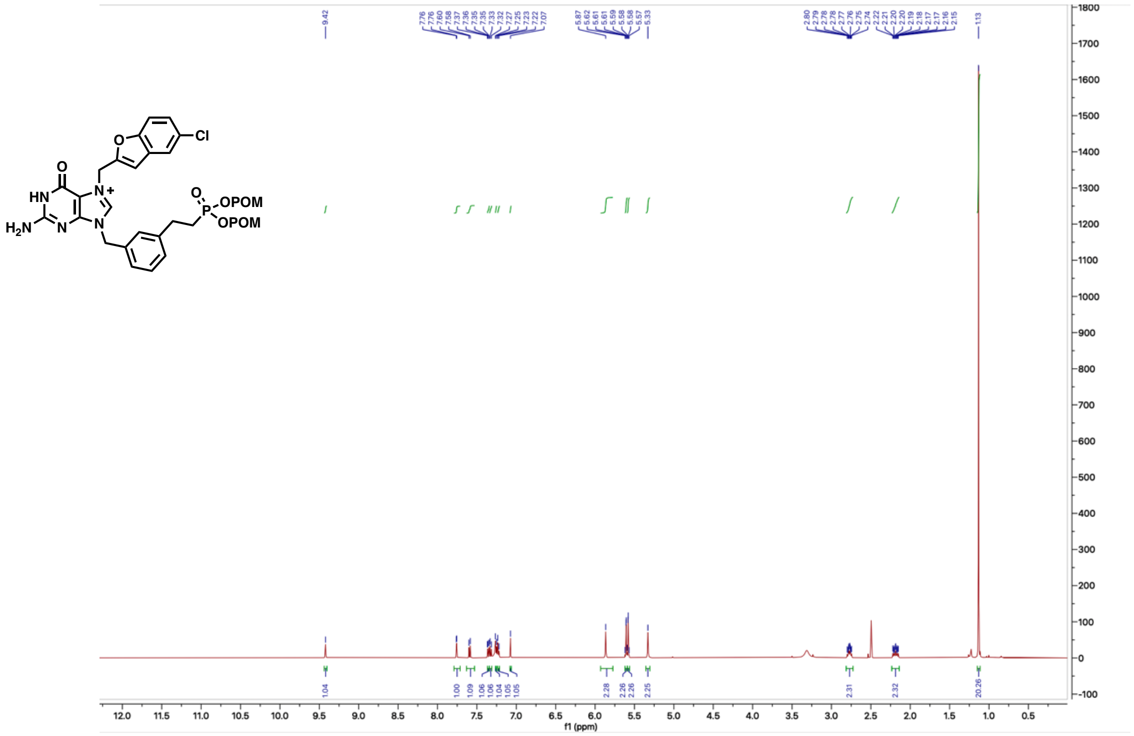

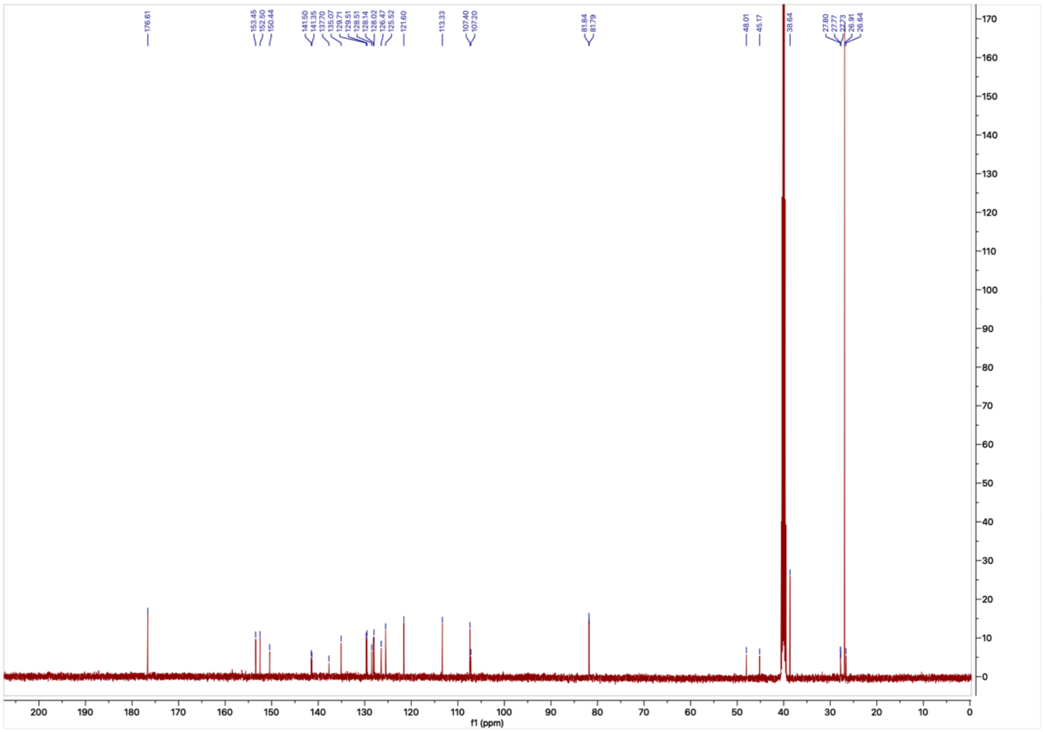

### Compound 6

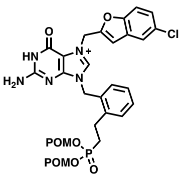


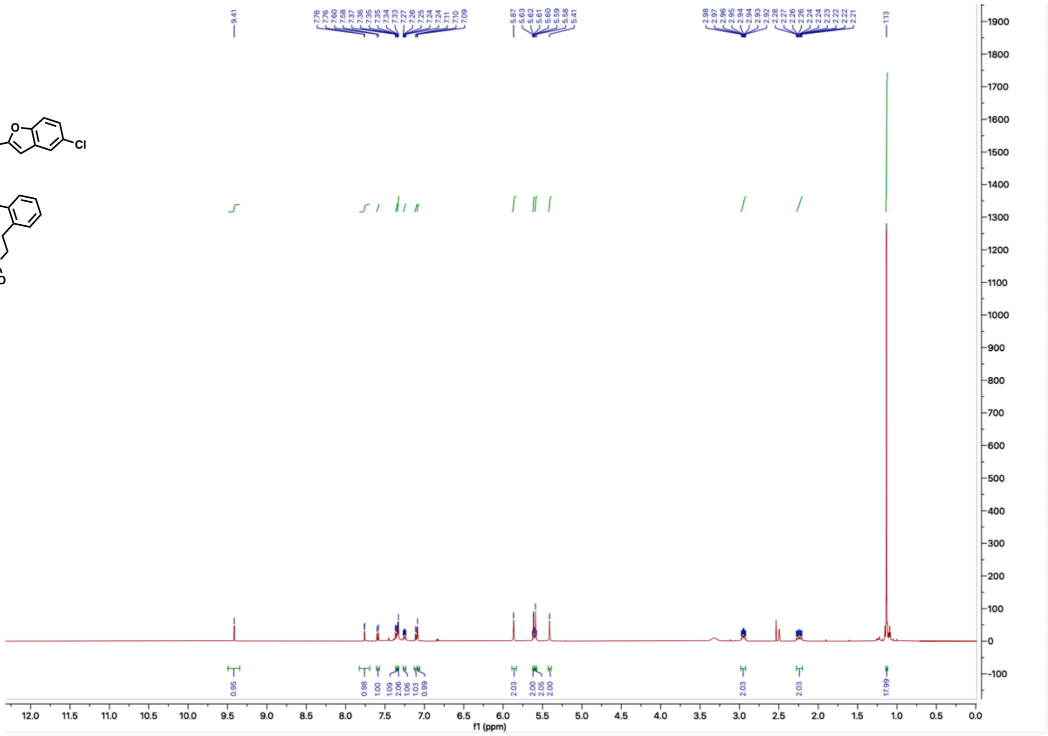


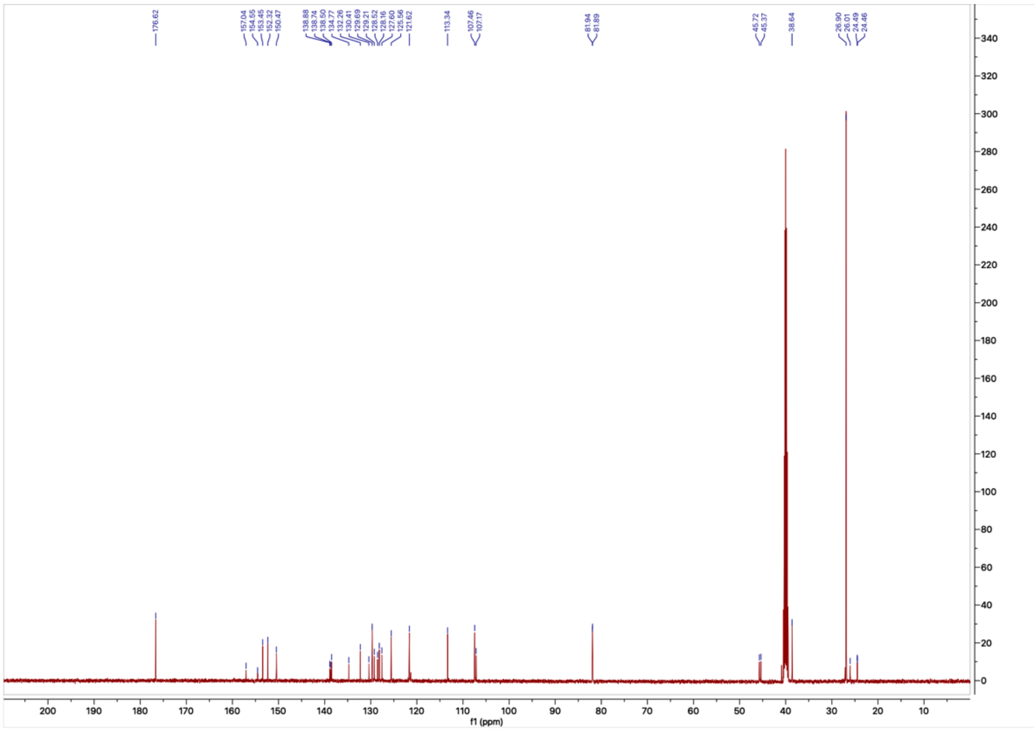

#### Compound 7

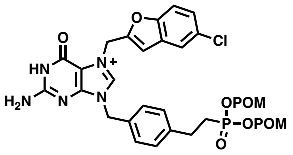


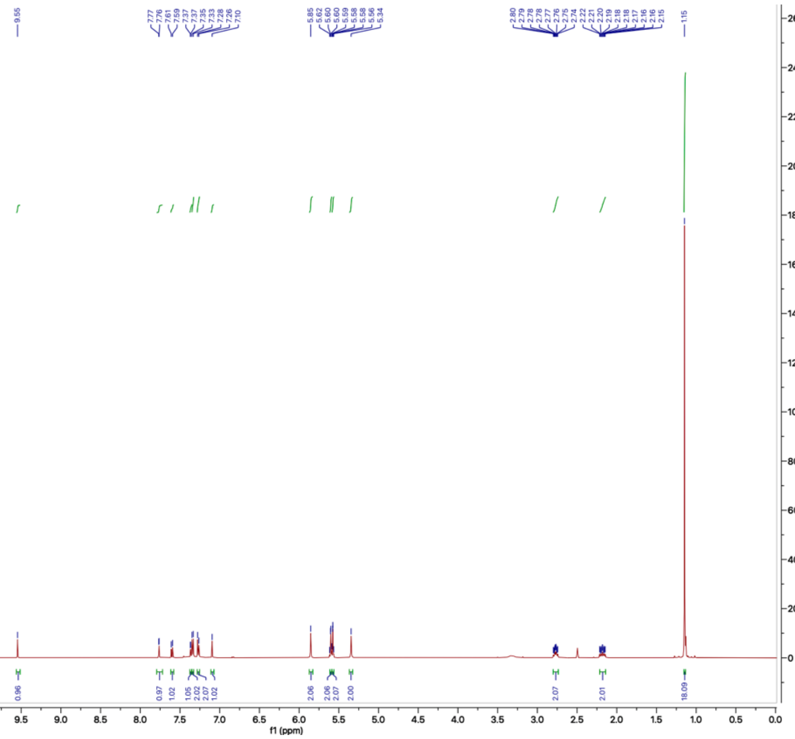


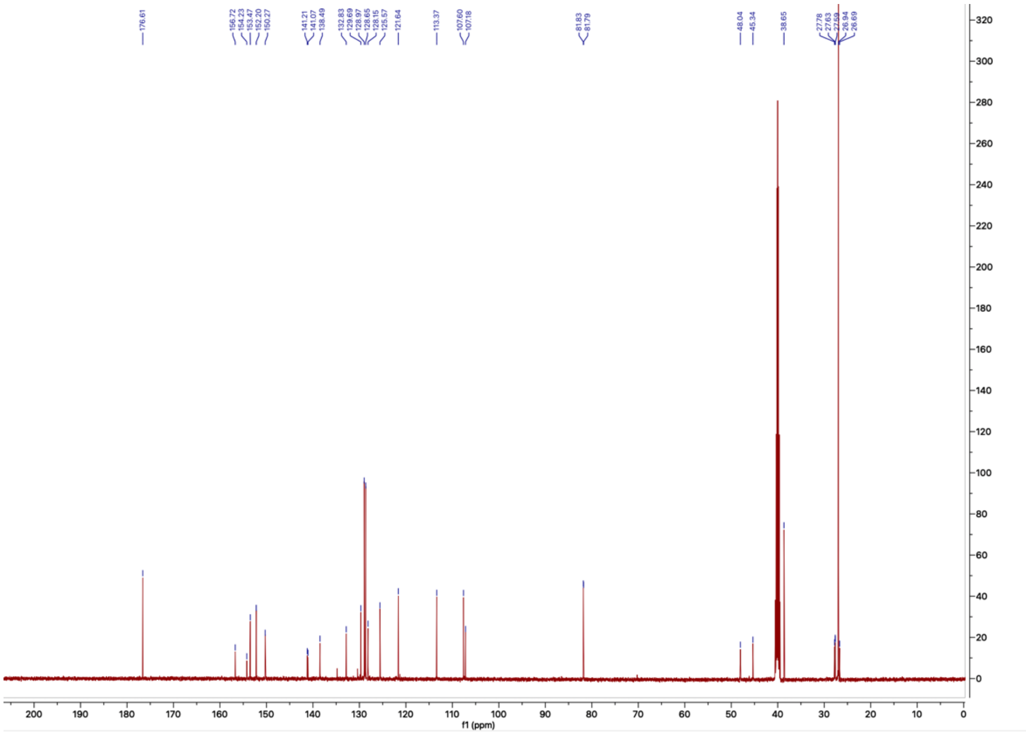

#### B. General Methods

**General Chemistry Materials and Methods.** All purchased solvents and reagents were used without further purification. Reactions were monitored by thin-layer chromatography (TLC) carried out on 0.25-mm SiliCycle silica gel plates (60F-254) using UV-light (254 nm). Flash chromatography was performed using SiliaFlash P60 silica gel. Analytical RP-HPLC was performed using an Agilent 1260 Infinity HPLC equipped with an ZORBAX Eclipse XDB-C18 colun (4.6 x 150 mm; 5 μm) with detection at 254 nm. Semi-preparative HPLC was carried out on Agilent 1260 Infinity HPLC equipped with a PrepHT XDB-C18 column (21.2 x 150 mm; 5 μm) with detection at 254 nm). General Method A: 30−60% MeCN/$H_2O$ (0.1% FA) over 30 min at 12 mL/min. General Method B: 5−80% MeCN/$H_2O$ (0.1% FA) over 30 min at 12 mL/min. NMR spectra were performed on a 300 MHz Bruker and 400 MHz Bruker instrument calibrated using a solvent peak as an internal reference. Chemical shifts ($\delta$ values) are reported in parts per million and are referenced to the deuterated residual solvent peak. Mass spectrometry (MS) was performed using an Agilent 6230 TOF LC/MS spectrometer using ESI ionization with an accuracy of 2 ppm. All compounds were found to be >95% pure by HPLC analysis.

**General Biology Materials.** EDA-m7GTP-5-FAM (NU-824-5FM) was purchased from Jena Bioscience and used as received. The following antibodies were used in this study: eIF4E (Cell Signaling Technology, 9742), eIF4G (Cell Signaling Technology, 2498), actin-HRP (Santa Cruz Biotechnology, sc-47778), c-Myc (Cell Signaling Technology, 13987), ODC1 (Novus Biologicals, 2878R), cyclin D1 (Cell Signaling Technology, 2978) and cyclin D3 (Cell Signaling Technology, 2936).

**General Cell Culture.** A2058 and A375 cells were grown in DMEM, 10% FBS, and 2 mM L-Glutamine at 37 °C with 5% $CO_2$ in a humidified incubator. SKMEL2 cells were grown in MEM, 10% FBS, 2 mM L-Glutamine, 1% sodium pyruvate, and 1% non-essential amino acids (NEAA) at 37 °C with 5% $CO_2$ in a humidified incubator.

**Cell Viability Assay.** The Cell Titer-Glo® assay kit was purchased from Promega and performed according to the manufacturer's instructions. Briefly, 2,500 A2058 cells were plated in a white, 96-well tissue culture-treated plate. SKMEL2 and A375 cells were seeded at 4,000 cells per well. Cells were treated with the compounds at different concentrations in quadruplicate and incubated for 48 hours. After 48 h, the cell culture media was replaced with 70 µl of OptiMEM and then lysed with 70 µl of Cell Titer-Glo® reagent. Total luminescence was read within 1 h using a BioTek Cytation 3 reader. Data was normalized and processed in GraphPad Prism.

**Total Cell Lysate and Western Blot.** A2058 cells were grown in a 6 well plate to be 50% confluent the next day (about 140,000 cells per well). After 16–24 h, cells were treated with 45 µM of compound (the $EC_{50}$ determined from dose-response studies) or DMSO control and incubated for 6 h. After 6 h, cells were harvested in 200–350 µL of RIPA buffer (10 mM Tris-HCl, 150 mM NaCl, 1% Triton, 1% sodium deoxycholate, 0.1% SDS, pH 7.2). Total protein was quantified using the BCA assay and analyzed by Western blot.

**Data and Statistical Analysis.** All data was analyzed using GraphPad Prism version 9.5.1 for Mac OS (GraphPad Software, www.graphpad.com). Graphs show mean ± standard deviation as described in the figure legends.

**CETSA and Western Blot.** Cultured HeLa were trypsinized and washed with PBS. Cell pellets were then diluted in PBS supplemented with protease inhibitors. Cell suspensions were freeze-thawed 3× using liquid nitrogen. The soluble fraction (lysate) was separated from the cell debris by centrifugation at 15,000 rpm for 25 min at 4°C. Cell lysates (150 µg total protein) were then divided and treated with either control (water) or compounds **7** (100 µM). After 1 h incubation at room temperature, lysates were divided into 90-µL aliquots and heated at 42 °C, 49 °C, or 52 °C for 3 min followed by cooling at 20 °C for 3 min. Heated lysates were centrifuged at 15,000 rpm for 25 min at 4°C to separate the soluble fractions from precipitates. Supernatants were transferred to new tubes and analyzed by SDS-PAGE. Samples were run on a 4−12% Bis-Tris gel and transferred to PVDF membrane in Towbin's Buffer. The membrane was blocked in 5% milk for 1 h at 25 °C, and then incubated with a primary antibody (overnight at 4 °C) and secondary antibody (1 h at 25 °C). Proteins were visualized using a Biorad ChemiDoc imaging system.

**Densitometry.** Scanned Western blot images were processed using ImageJ software. The pixels in each band gave a raw reading. The baseline was subtracted from each raw reading. Raw readings were normalized to actin, and then ratios of treatment to control were calculated. Graphs were plotted in GraphPad Prism.

**Protein Expression and Purification.** Human eIF4E subcloned into a pET19b vector modified to include a 10× His tag and a PreScission protease cut-site between the tag and the beginning of the eIF4E gene was used for expression.[1] pET19bpp-eIF4E plasmid was transformed into BL21(DE3) cells. Cells were grown at 37 °C to an $OD_{600}$ of 0.6–0.8, induced with 1 mM IPTG, and grown for 16 h at 18 °C. The cells were pelleted and lysed through sonication in lysis buffer (50 mM Tris-HCl, pH 8, 500 mM NaCl, 50 mM imidazole, 20 mM β-mercaptoethanol, 5 mM DTT, and 1% Tween-20 with protease inhibitors). The cell lysate was centrifuged at 38,000g for 2 h and then incubated with Ni-NTA resin for 1 h at 4 °C. The resin was washed with 15 mL of lysis buffer (3−4×, 15 min each, in 50 mM Tris-HCl, pH 8, 500 mM NaCl, 25 mM imidazole, 5 mM DTT). eIF4E was eluted with 5-mL aliquots (total ~ 25 mL) of elution buffer (10 mM Tris-HCl, pH 8, 500 mM NaCl, 100 mM imidazole, 5 mM DTT). Protein was then dialyzed overnight at 4 °C in dialysis buffer (20 mM Tris buffer, pH 7.4, 100 mM NaCl, 2 mM DTT). After dialysis, protein was concentrated and purified using a FPLC Superdex 75 16/10 preparative column. Pure protein was aliquoted and stored in −80 °C. The yield was ~1 mg from 1 L of cell culture.

**FP Assay.** FP measurements were performed on a Biotek Cytation 3 microplate reader equipped with excitation (485 ± 20 nm) and emission (528 ± 20 nm) polarization filters. Experiments were carried out at room temperature in 384-well non-binding low volume black microplates with sample volume of 18 μL per well. Each condition was done in quadruplicate and each experiment in triplicate. eIF4E FP buffer was 50 mM HEPES (pH 7.2), 100 mM KCl, 0.5 mM EDTA, 1 mM DTT, 0.0025% Tween 20 and 0.05 mg/mL BSA. To determine $K_d$ values, 15 nM EDA-m7GTP-5-FAM was mixed with increasing concentration of His10-eIF4E (0−2 μM). Before FP measurements, the plate was shaken at 150 rpm at room temperature for 40 min. For competition experiments, a mixture containing 15 nM probe and 95 nM protein in FP buffer was incubated with the increasing concentrations of compound **7** (0−80 μM). Plates were shaken for 40 min at room temperature and FP was measured. Values were plotted using GraphPad Prism.

**eIF4E expression and purification for crystallography.** The gene encoding eIF4E (residues 27-217) was cloned into a pET19b expression vector and transformed into Rosetta2 cells. The protein was expressed and purified as described in Papadopoulos *et al.*[2] with the following modification that was referenced in Cárdenas *et al.*[3] Eluant from the diethylaminoethylcellolose column was incubated with the cap resin overnight and then eluted the following day. After each elution, the protein was buffer exchanged into 10 mM HEPES, pH 7.5, 125 mM NaCl, and 1 mM TCEP then concentrated to 0.5 to 1.0 mg/mL and flash frozen in liquid nitrogen.

**Crystallization and Structure Determination.** Crystals of the eIF4E: **5-PA** complex were achieved from soaking experiments. eIF4E protein was concentrated to 4.6 mg/mL and native crystals were grown in well solution containing 16–26% Peg 3350, 0.1 M MES pH 6.0, 10% isopropanol and 2 mM $CaCl_2$. A powdered form of **5-PA** was added directly to the drop containing the crystals and incubated overnight. The following day, the crystals were cryoprotected in 26% Peg 3350, 0.1 M MES pH 6.0, 2 mM $CaCl_2$ and 20% glycerol and flash frozen for data collection. Diffraction data were collected at the Advanced Photon Source beamline 21-ID-D at Argonne National Laboratory and processed with HKL2000.[4] The structure was solved by molecular replacement in Phaser[5] using the protein component of 4TQC as a search model. Iterative rounds of model building and refinement were completed using Coot[6] and Buster[7], respectively. Coordinates and geometric restraints for the ligand and inhibitor were created using Grade.[7]

eIF4E crystals grew in space group P2 with 2 molecules in the asymmetric unit. After molecular replacement, difference electron density corresponding to **5-PA** was observed in the binding site of the B chain, while density for $m^7$GTP was observed in the A chain. In chain A, there was clear electron density for residues 27−217 with the exception of residues 208−211. In the B chain, residues 27−28, 83−88, 120−124, and 209−210 were disordered. The structure was validated using Molprobity[8] and ligand statistics were obtained from the PDB validation server. Data collection and refinement statistics are given in Table S2.

#### C. Synthetic Methods

**Scheme 1.** Synthesis of **2**

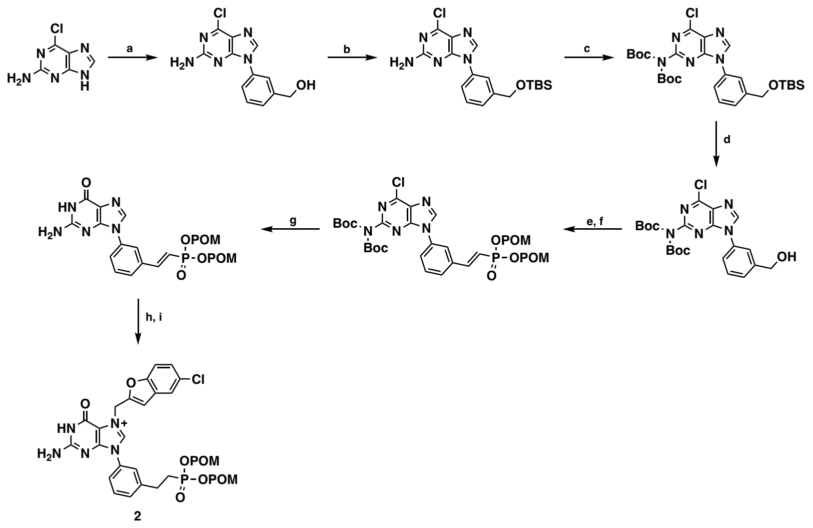

Reagents and conditions: (a) 3-(hydroxymethyl)phenyl boronic acid, 1,10-phenanthroline, $Cu(OAc)_2$, DMF, 23 °C; (b) TBSCl, imidazole, DMF, 23 °C; (c) $Boc_2O$, DMAP, THF, 23 °C; (d) TBAF (1M in THF), THF, 0 °C; (e) DMP, $CH_2Cl_2$, 23 °C; (f) NaH, tetraPOM methylene diphosphonate, THF, 0 °C to 23 °C; (g) $CH_2O_2$:$H_2O$ (1:1), 40 °C; (h) 10% Pd/C, $H_2$, MeOH:$H_2O$ (4:1), 23 °C; (i) 2-(bromomethyl)-5-chlorobenzofuran, DMSO, 50 °C.

**2**, step a: *(3-(2-amino-6-chloro-9H-purin-9-yl)phenyl)methanol.* 3-(hydroxymethyl)phenyl boronic acid (1.79 g, 11.79 mmol) was diluted in DMF (0.2 M, 30 mL) then treated with 1,10-phenanthroline (2.12 g, 11.79 mmol), $Cu(OAc)_2$ (2.14 g, 11.79 mmol) and 2-amino-6-chloropurine at 23 °C. After 4 d the reaction mixture was concentrated then diluted in EtOAc (200 mL) and washed with an aqueous solution of EDTA (1.0 g in 150 mL of $H_2O$) and brine. The organic layer was dried over $Na_2SO_4$ then purified by column chromatography (2–4% MeOH/$CH_2Cl_2$) to provide 3-(hydroxymethyl)phenyl purine (636 mg, 40%). $^1H$ NMR (300 MHz, DMSO-$d_6$) δ 8.49 (s, 1H), 7.68 (d, $J$ = 10.5 Hz, 2H), 7.54 (t, $J$ = 7.5 Hz, 1H), 7.42 (d, $J$ = 7.7 Hz, 1H), 7.04 (s, 1H), 5.38 (t, $J$ = 5.6 Hz, 1H), 4.59 (d, $J$ = 5.6 Hz, 2H).

**2**, step b: *9-(3-(((tert-butyldimethylsilyl)oxy)methyl)phenyl)-6-chloro-9H-purin-2-amine.* Purine from step a (100 mg, 0.36 mmol) was diluted in DMF (0.2 M, 2 mL) then treated with TBSCl (82 mg, 0.91 mmol) and imidazole (61 mg, 0.55 mmol) at 23 °C. After 16 h the reaction mixture was quenched with $H_2O$ (5 mL) then diluted in EtOAc (150 mL) and washed with $H_2O$ (x2, 50 mL) and brine (100 mL). The isolated organic layer was dried over $Na_2SO_4$ then concentrated to dryness and purified by column chromatography (40-60% ethyl acetate/hexanes) to yield the product (166 mg, >99%). $^1$H NMR (300 MHz, MeOD-d$_4$) δ 8.40 (s, 1H), 7.75 (s, 1H), 7.67 (d, $J$ = 7.4 Hz, 1H), 7.56 (t, $J$ = 8.0 Hz, 1H), 7.48 (d, $J$ = 8.0 Hz, 1H), 4.88 (s, 2H), 0.98 (s, 9H), 0.16 (s, 6H).

**2**, step c: *tert-butyl (tert-butoxycarbonyl)(9-(3-(((tert-butyldimethylsilyl)oxy)methyl)phenyl)-6-chloro-9H-purin-2-yl)carbamate*. Purine from step b was diluted in THF (0.2 M, 2 mL) then treated with $Boc_2O$ (236 mg, 1.01 mmol) and DMAP (5 mg, 0.04 mmol) at 23 °C. After 4 d the reaction mixture was quenched with $NH_4Cl_{(sat)}$ (5 mL) then diluted in EtOAc (150 mL) and washed with $NH_4Cl_{(sat)}$ (x2, 50 mL) and brine (50 mL). The isolated organic layer was dried over $Na_2SO_4$ then concentrated and purified by column chromatography (25% ethyl acetate/hexanes) to yield the product (200 mg, 95%). $^1$H NMR (300 MHz, $CDCl_3$) δ 8.44 (s, 1H), 7.65 (s, 1H), 7.57 (t, $J$ = 6.8 Hz, 1H), 7.51 (d, $J$ = 7.8 Hz, 1H), 7.43 (d, $J$ = 7.7 Hz, 1H), 4.83 (s, 2H), 1.43 (s, 18H), 0.94 (s, 9H), 0.12 (s, 6H).

**2**, step d: *tert-butyl (tert-butoxycarbonyl)(6-chloro-9-(3-(hydroxymethyl)phenyl)-9H-purin-2-yl)carbamate*. Purine from step c (180 mg, 0.32 mmol) was diluted in THF (0.15 M, 3 mL) then treated with a solution of TBAF (1 M in THF, 350 µL) at 0 ºC. After 1 h the reaction mixture was quenched with $H_2O$ (5 mL) then diluted further in EtOAc (100 mL) and washed with $H_2O$ (50 mL) and brine (50 mL). The isolated organic layer was dried over $Na_2SO_4$ then concentrated to dryness and purified by column chromatography (40–60% ethyl acetate/hexanes) to provide the product (120 mg, 79%). $^1$H NMR (300 MHz, MeOD-d$_4$) δ 8.93 (s, 1H), 7.83 (s, 1H), 7.71 (d, $J$ = 7.8 Hz, 1H), 7.61 (t, $J$ = 7.8 Hz, 1H), 7.54 (d, $J$ = 7.5 Hz, 1H), 4.75 (s, 2H), 1.44 (s, 18H).

**2**, steps e and f: *(E)-(((3-(2-(bis(tert-butoxycarbonyl)amino)-6-chloro-9H-purin-9-yl)styryl)phosphoryl)bis(oxy))bis(methylene) bis(2,2-dimethylpropanoate)*. Purine from step d (200 mg, 0.42 mmol) was diluted in $CH_2Cl_2$ (0.12 M, 5 mL) then treated with Dess-Martin Periodinane (233 mg, 0.55 mmol) at 23 °C. After 1 h the reaction mixture was quenched with $Na_2S_2O_{3(sat)}$ (5 mL) then diluted further with $CH_2Cl_2$ (100 mL) and washed with $NaHCO_{3(sat)}$ (x2, 50 mL) and brine (x2, 50 mL). The isolated organic layer was dried over $Na_2SO_4$ then concentrated to dryness to provide the crude aldehyde which was used without further purification. TetraPOM methylenediphosphonate (398 mg, 0.63 mmol) was diluted in THF (0.07 M, 6 mL) then treated with NaH (60% in mineral oil, 50 mg, 1.26 mmol) at 0 °C. After 5 min the reaction mixture was treated dropwise with the crude aldehyde in a minimal amount of THF (~1 mL) at 0 °C. After 1 h the reaction mixture was removed from the cooling bath and allowed to reach 23 °C. After 3 h the reaction mixture was quenched with $NH_4Cl_{(sat)}$ (5 mL) then diluted with EtOAc (100 mL) and washed with $NH_4Cl_{(sat)}$ (x2, 50 mL) and brine (50 mL). The isolated organic layer was dried over $Na_2SO_4$ then concentrated to dryness and purified by column chromatography (30–50% ethyl acetate/hexanes) to provide the vinyl phosphonate product (262 mg, 80% over 2 steps). $^1$H NMR (300 MHz, MeOD-d$_4$) δ 8.99 (s, 1H), 8.15 (s, 1H), 7.93 (d, $J$ = 7.9 Hz, 1H), 7.82 (d, $J$ = 6.8 Hz, 1H), 7.74 (d, $J$ = 8.0 Hz, 1H), 7.70 (m, 1H), 6.70 (t, $J$ = 18.2 Hz, 1H), 5.75 (d, $J$ = 13.1 Hz, 4H), 1.46 (s, 18H), 1.21 (s, 18H).

**2**, step g: *(E)-(((3-(2-amino-6-oxo-1,6-dihydro-9H-purin-9-yl)styryl)phosphoryl)bis(oxy))bis(methylene) bis(2,2-dimethylpropanoate)*. Purine from step f (250 mg, 0.32 mmol) was diluted in an aqueous solution of formic acid (1:1, 0.06 M, 6 mL) and warmed to 40 °C. After 24 h the reaction mixture was cooled to 23 °C then concentrated to dryness. The crude residue was purified by column chromatography (2–10% MeOH/$CH_2Cl_2$) to yield the product (87 mg, 49%). $^1$H NMR (400 MHz, MeOD-d$_4$) δ 8.01 (s, 1H), 7.62 (s, 1H), 7.58 (d, $J$ = 8.0 Hz, 1H), 7.49 (t, $J$ = 5.6 Hz, 1H), 7.35 (d, $J$ = 7.5 Hz, 1H), 5.69 (dq, $J$ = 12.8, 1.6 Hz, 4H), 3.00 (m, 2H), 2.33 (m, 2H), 1.24 (s, 18H).

**2**, steps h and i: *(((3-(2-amino-6-oxo-1,6-dihydro-9H-purin-9-yl)phenethyl)phosphoryl)bis(oxy))bis(methylene) bis(2,2-dimethylpropanoate) (**2**)*. Guanine analog **7** (40 mg, 0.07 mmol) was diluted in an aqueous solution of MeOH (4:1, 0.01 M, 10 mL) then treated with 10% Pd/C (10 mg, 20% w/w) and the resulting mixture was back-filled with a balloon of $H_{2(g)}$ at 23 °C. After 24 h the reaction mixture was filtered through a pad of celite then the filtrate was concentrated to provide the guanine analog **8** (37 mg, >99%) which was used without further purification. The product (30 mg, 0.05 mmol) was diluted in DMSO (0.06 M, 1 mL) then treated with 2-(bromomethyl)-5-chlorobenzofuran (40 mg, 0.16 mmol) at 50 °C. After 48 h the reaction mixture was loaded directly on to silica gel and purified by column chromatography (2–10% MeOH/$CH_2Cl_2$) then purified further by RP-HPLC Method A to provide cap analogue prodrug **2** (12 mg, 32%).

**Scheme 2.** Synthesis of **3**

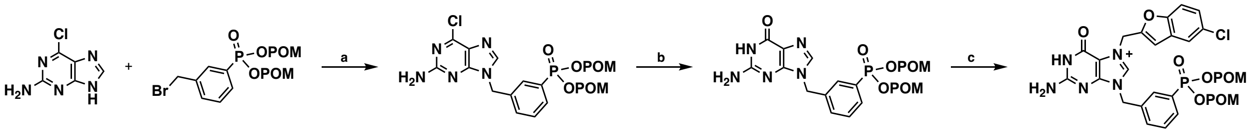


3

Reagents and conditions: (a) bis-POM benzyl bromide, $Cs_2CO_3$, DMF, 23 °C; (b) $CH_2O_2$:$H_2O$ (1:1), 40 °C; (c) 2-(bromomethyl)-5-chlorobenzofuran, DMSO, 50 °C.

**3**, step a: *(((3-((2-amino-6-chloro-9H-purin-9-yl)methyl)phenyl)phosphoryl)bis(oxy))bis(methylene) bis(2,2-dimethylpropanoate)*. 2-amino-6-chloropurine (70 mg, 0.42 mmol) was diluted in DMF (0.4 M, 1 mL) then treated with bis-POM benzyl bromide (200 mg, 0.42 mmol) and $Cs_2CO_3$ (136 mg, 0.42 mmol) at 23 °C. After 20 h the reaction mixture was cooled to 23 °C then diluted further with EtOAc (200 mL) and washed with $H_2O$ (50 mL) and brine (x2, 50 mL). The isolated organic layer was dried over $Na_2SO_4$ then concentrated to dryness and purified by column chromatography (60–80% ethyl acetate/hexanes) to yield the product (32 mg, 14%). $^1$H NMR (400 MHz, MeOD-d$_4$) δ 8.19 (s, 1H), 7.92 (d, $J$ = 14.7 Hz, 1H) 7.73 (m, 2H), 7.55 (m, 1H), 5.73 (d, $J$ = 14.1 Hz, 4H), 5.40 (s, 2H), 1.02 (s, 18H).

**3**, step b: *(((3-((2-amino-6-oxo-1,6-dihydro-9H-purin-9-yl)methyl)phenyl)phosphoryl)bis(oxy))bis(methylene) bis(2,2-dimethylpropanoate)*. Purine from step a (30 mg, 0.052 mmol) was diluted in an aqueous solution of formic acid (1:1, 0.06 M, 1 mL) then warmed to 40 °C. After 21 h the reaction mixture was concentrated to dryness then purified by column chromatography (4–8% $MeOH/CH_2Cl_2$) to yield the product (15 mg, 53%). $^1$H NMR (400 MHz, MeOD-$d_4$) δ 7.89 (dt, $J$ = 14.8, 1.7 Hz, 1H), 7.83 (s, 1H), 7.75 (tt, $J$ = 7.5, 1.3 Hz, 1H), 7.70 (dq, $J$ = 6.2, 1.3 Hz), 7.56 (m, 1H), 5.75 (d, 1.7 Hz, 4 H), 5.33 (s, 2H), 1.05 (s, 18H).

**3**, step c: *2-amino-9-(3-(bis((pivaloyloxy)methoxy)phosphoryl)benzyl)-7-((5-chlorobenzofuran-2-yl)methyl)-6-oxo-6,9-dihydro-1H-purin-7-ium (**3**)*. Purine from step b (15 mg, 0.027 mmol) was diluted in DMSO (0.06 M, 0.5 mL) then treated with 2-(bromomethyl)-5-chlorobenzofuran (20 mg, 0.082 mmol) and warmed to 50 °C. After 48 h the reaction mixture was loaded directly on to silica gel then purified by column chromatography (5–12% $MeOH/CH_2Cl_2$) and further purified by RP-HPLC Method A to yield the cap analogue prodrug **3** (4 mg, 21%).

**Scheme 3.** Synthesis of **4**

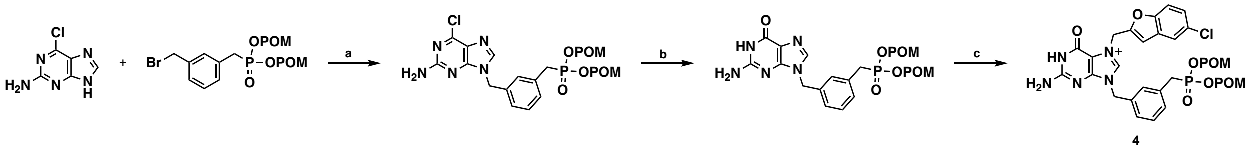


Reagents and conditions: (a) bis-POM benzyl bromide, $Cs_2CO_3$, DMF, 23 °C; (b) $CH_2O_2$:$H_2O$ (1:1), 40 °C; (c) 2-(bromomethyl)-5-chlorobenzofuran, DMSO, 50 °C.

**4**, step a: *(((3-((2-amino-6-chloro-9H-purin-9-yl)methyl)benzyl)phosphoryl)bis(oxy))bis(methylene) bis(2,2-dimethylpropanoate)*. 2-amino-6-chloropurine (308 mg, 1.82 mmol) was diluted in DMF (0.2 M, 10 mL) then treated with bis-POM benzyl bromide (900 mg, 1.82 mmol) and $K_2CO_3$ (300 mg, 2.18 mmol) and warmed to 60 °C. After 19 h the reaction mixture was concentrated then diluted further with EtOAc (150 mL) and washed with $H_2O$ (x2, 50 mL) and brine (x2, 50 mL). The isolated organic layer was dried over $Na_2SO_4$ then concentrated to dryness and purified by column chromatography (60–80% ethyl acetate/hexanes) to yield the product (96 mg, 10%). $^1$H NMR (400 MHz, MeOD-$d_4$) δ 8.13 (s, 1H), 7.34 (s, 1H), 7.27 (m, 3H), 5.59 (dq, $J$ = 13.0, 3.2 Hz, 4H), 5.32 (s, 2H), 3.34 (d, $J$ = 22.2 Hz, 2H), 1.18 (s, 18H).

**4**, step b: *(((3-((2-amino-6-oxo-1,6-dihydro-9H-purin-9-yl)methyl)benzyl)phosphoryl)bis(oxy))bis(methylene) bis(2,2-dimethylpropanoate)*. Purine from step a (70 mg, 0.12 mmol) was diluted in an aqueous solution of formic acid (1:1, 0.06 M, 2 mL) then warmed to 40 °C. After 22 h the reaction mixture was then concentrated to dryness and purified by column chromatography (2–10% $MeOH/CH_2Cl_2$) to yield the product (36 mg, 54%). $^1$H NMR (400 MHz, MeOD-$d_4$) δ 7.80 (bs, 1H), 7.32 (m, 2H), 7.25 (m, 2H), 5.59 (dq, $J$ = 12.9, 4.9 Hz, 4H), 5.25 (s, 2H), (d, $J$ = 19.7 Hz, 2H), 1.20 (s, 18H).

**4**, step c: *2-amino-9-(3-((bis((pivaloyloxy)methoxy)phosphoryl)methyl)benzyl)-7-((5-chlorobenzofuran-2-yl)methyl)-6-oxo-6,9-dihydro-1H-purin-7-ium (**4**)*. Purine from step b (30 mg, 0.053 mmol) was diluted in DMSO (0.06 M, 1 mL) then treated with 2-(bromomethyl)-5-chlorobenzofuran (39 mg, 0.16 mmol) and warmed to 50 °C. After 48 h the reaction mixture was loaded directly onto silica gel then purified by column chromatography (4–8% $MeOH/CH_2Cl_2$) and further purified by RP-HPLC Method A to yield cap analogue prodrug **4** (8 mg, 21%).

**Scheme 4.** Synthesis of **5**

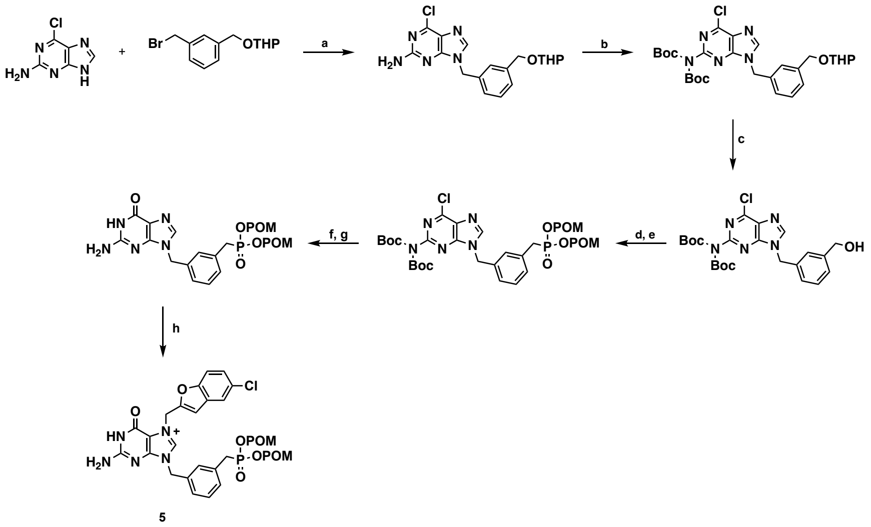

Reagents and conditions: (a) benzyl bromide, $K_2CO_3$, DMF, 80 °C; (b) $Boc_2O$, DMAP, THF, 23 °C; (c) pTSA monohydrate, MeOH, 23 °C; (d) DMP, $CH_2Cl_2$, 23 °C; (e) NaH, tetraPOM methylenediphosphonate, THF, 0 °C to 23 °C; (f) $CH_2O_2$:$H_2O$ (1:1), 40 °C; (g) 10% Pd/C, $H_2$, MeOH:$H_2O$ (4:1), 23 °C; (h) 2-(bromomethyl)-5-chlorobenzofuran, DMSO, 50 °C.

**5**, step a: *6-chloro-9-(3-(((tetrahydro-2H-pyran-2-yl)oxy)methyl)benzyl)-9H-purin-2-amine*. 2-amino-6-chloropurine (1.25 g, 7.4 mmol) was diluted in DMF (0.2 M, 40 mL) then treated with 2-((3-(bromomethyl)benzyl)oxy)tetrahydro-2*H*-pyran (2.1 g, 7.4 mmol) and $K_2CO_3$ (3.0 g, 22.2 mmol) at 80 °C. After 21 h the reaction mixture was concentrated to dryness then diluted in EtOAc (300 mL) and washed with $H_2O$ (100 mL) and brine (x3, 100 mL). The isolated organic layer was dried over $Na_2SO_4$ then concentrated to dryness and purified by column chromatography to provide the product (1.28 g, 46%). $^1$H NMR (400 MHz, $CDCl_3$) δ 8.12 (s, 1H), 7.30 (m, 3H), 5.35 (s, 2H), 4.70 (d, $J$ = 12.2 Hz, 1H), 4.65 (t, $J$ = 3.9 Hz, 1H), 4.48 (d, $J$ = 12.2 Hz, 1H), 3.82 (m, 1H), 3.48 (m, 1H), 1.79 (m, 1H), 1.68 (m, 1H), 1.54 (m, 4H).

**5**, step b: *tert-butyl (tert-butoxycarbonyl)(6-chloro-9-(3-(((tetrahydro-2H-pyran-2-yl)oxy)methyl)benzyl)-9H-purin-2-yl)carbamate*. Purine from step a (1.2 g, 3.21 mmol) was diluted in THF (0.1 M, 35 mL) then treated with $Boc_2O$ (2.1 g, 9.64 mmol) and DMAP (40 mg, 0.32 mmol) at 23 °C. After 2 d the reaction mixture was concentrated then diluted in EtOAc (200 mL) and washed with $H_2O$ (100 mL) and brine (100 mL). The isolated organic layer was dried over $Na_2SO_4$ then concentrated to dryness and purified by column chromatography (1–2% MeOH/$CH_2Cl_2$) to yield the Boc-protected product (1.58 g, 93%). $^1$H NMR (400 MHz, MeOD-$d_4$) δ 8.73 (s, 1H), 7.36 (m, 1H), 7.34 (m, 1H), 7.29 (m, 1H), 5.67 (s, 2H), 4.73 (d, $J$ = 12.1 Hz, 1H), 4.67 (t, $J$ = 4.3 Hz, 1H), 4.49 (d, $J$ = 12.2 Hz, 1H), 3.86 (m, 1H), 3.51 (m, 1H), 1.82 (m, 1H), 1.73 (m, 1H), 1.59 (m, 4H), 1.38 (s, 18H).

**5**, step c: *tert-butyl (tert-butoxycarbonyl)(6-chloro-9-(3-(hydroxymethyl)benzyl)-9H-purin-2-yl)carbamate*. Boc-protected purine from step b (1.5 g, 2.81 mmol) was diluted in MeOH (0.1 M, 30 mL) then treated with pTSA monohydrate (53 mg, 0.28 mmol) at 23 ºC. After 1 h the reaction mixture was concentrated then diluted in $CH_2Cl_2$ (200 mL) and washed with $NaHCO_{3(sat)}$ (x2, 100 mL), $H_2O$ (100 mL) and brine (100 mL). The isolated organic layer was dried over $Na_2SO_4$ then concentrated to dryness and purified by column chromatography (30–60% ethyl acetate/hexanes) to provide the product (1.26 g, 92%). $^1$H NMR (400 MHz, MeOD-$d_4$) δ 8.71 (s, 1H), 7.38 (s, 1H), 7.34 (m, 1H), 7.33 (d, $J$ = 1.1 Hz, 1H), 7.26 (m, 1H), 5.56 (s, 2H), 4.59 (s, 2H), 1.39 (s, 18H).

**5**, steps d and e: *(E)-(((3-((2-(bis(tert-butoxycarbonyl)amino)-6-chloro-9H-purin-9-yl)methyl)styryl)phosphoryl)bis(oxy))bis(methylene) bis(2,2-dimethylpropanoate)*. Purine from step c (800 mg, 1.63 mmol) was diluted in $CH_2Cl_2$ (0.12 M, 15 mL) then treated with Dess-Martin Periodinane (902 mg, 2.12 mmol) at 23 °C. After 1 h the reaction mixture was quenched with $Na_2S_2O_{3(sat)}$ (5 mL) then stirred for an additional 30 min. The resulting mixture was diluted further with $CH_2Cl_2$ (200 mL) then washed with $H_2O$ (100 mL) and brine (100 mL). The isolated organic layer was dried over $Na_2SO_4$ then concentrated to dryness to furnish the crude aldehyde that was used without further purification. TetraPOM methylenediphosphonate (1.4 g, 2.24 mmol) was diluted in THF (0.06 M, 30 mL) then treated with NaH (60% in mineral oil, 200 mg, 4.8 mmol) at 0 °C. After 5 min the reaction mixture was treated with the crude aldehyde in a minimal amount of THF (5 mL) at 0 °C. After 1 h the reaction mixture was removed from the cooling bath and allowed to stir at 23 °C. After 2h the reaction mixture was quenched with $NH_4Cl_{(sat)}$ (10 mL) then concentrated and diluted further with EtOAc (200 mL). The resulting mixture was washed with $NH_4Cl_{(sat)}$ (100 mL) and brine (100 mL) then dried over $Na_2SO_4$ and concentrated to dryness. The resulting crude residue was purified by column chromatography (1% MeOH/$CH_2Cl_2$) to yield the vinyl phosphonate product (929 mg, 77% over 2 steps). $^1$H NMR (400 MHz, $CDCl_3$) δ 8.13 (s, 1H), 7.49 (q, $J$ = 8.5, 2 H), 7.41 (m, 2H), 7.33 (d, $J$ = 7.7 Hz, 1H), 6.32 (t, $J$ = 18.4 Hz, 1H), 5.73 (d, $J$ = 13.5 Hz, 4H), 5.46 (s, 2H), 1.45 (s, 18H), 1.20 (s, 18H).

**5**, step f and g: *(((3-((2-amino-6-oxo-1,6-dihydro-9H-purin-9-yl)methyl)phenethyl)phosphoryl)bis(oxy))bis(methylene) bis(2,2-dimethylpropanoate)*. Purine from step 3 (900 mg, 0.38 mmol) was diluted in an aqueous solution of formic acid (0.06 M, 8 mL) then warmed to 40 °C. After 22 h the reaction mixture was cooled to 23 °C then concentrated to dryness and the crude product was used without further purification. The crude guanine analogue was diluted in an aqueous solution of MeOH (4:1, 0.02 M, 50 mL) then treated with 10% Pd/C (130 mg, 20% w/w) and back-filled with $H_{2(g)}$ at 23 °C. After 24 h the reaction mixture was passed through a pad of celite then the filtrate was concentrated to dryness and purified by column chromatography (2–8% MeOH/$CH_2Cl_2$) to yield the product (489 mg, 77% over 2 steps). $^1$H NMR (400 MHz, DMSO-$d_6$) δ 10.60 (s, 1H), 7.73 (s, 1H), 7.27 (t, $J$ = 7.4 Hz, 1H), 7.17 (d, $J$ = 7.2 Hz, 1H), 7.13 (s, 1H), 7.05 (d, $J$ = 7.6 Hz, 1H), 6.49 (s, 2H), 5.60 (d, $J$ = 13.0 Hz, 4H), 5.14 (s, 2H), 2.75 (m, 2H), 2.17 (m, 2H), 1.16 (s, 18H).

**5**, step h: *2-amino-9-(3-(2-(bis((pivaloyloxy)methoxy)phosphoryl)ethyl)benzyl)-7-((5-chlorobenzofuran-2-yl)methyl)-6-oxo-6,9-dihydro-1H-purin-7-ium (**5**)*. Purine from step g (10 mg, 0.017 mmol) was diluted in DMSO (0.06 M, 0.30 mL) then treated with 2-(bromomethyl)-5-chlorobenzofuran (13 mg, 0.052 mmol) and warmed to 50 °C. After 48 h the reaction mixture was loaded directly on silica gel and purified by column chromatography (2–8% MeOH/$CH_2Cl_2$) and further purified by RP-HPLC Method A to yield the cap analogue prodrug **5** (5 mg, 42%).

**Scheme 5.** Synthesis of **6**

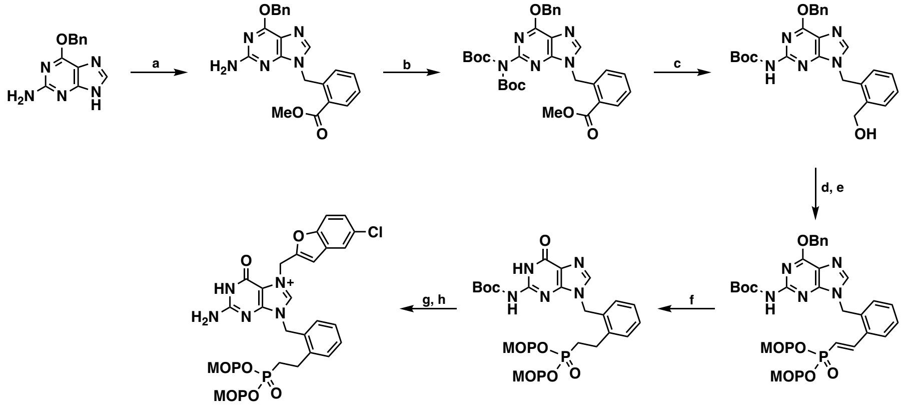


Reagents and conditions: (a) methyl 2-(bromomethyl)benzoate, $K_2CO_3$, DMF, 23 °C; (b) $Boc_2O$, DIPEA, DMAP, MeCN, 70 °C; (c) LAH, THF, 0 °C; (d) DMP, $NaHCO_3$, $CH_2Cl_2$, 23 °C; (e) NaH, TetraPOM methylenediphosphonate, THF, 0 °C to 23 °C; (f) 10% Pd/C, $H_2$, MeOH; (g) $CH_2Cl_2$:TFA (2:1), 23 °C; (h) 2-(bromomethyl)-5-chlorobenzofuran, DMSO, 50 °C.

**6**, step a: *Methyl 2-((2-amino-6-(benzyloxy)-9H-purin-9-yl)methyl)benzoate*. $O^6$-benzylguanine (500 mg, 2.07 mmol) was diluted in DMF (0.5 M, 30 mL) then treated with methyl 2-(bromomethyl)benzoate (566 mg, 2.48 mmol) and $K_2CO_3$ (1.43 g, 10.36 mmol) at 23 °C. After 18 h the reaction mixture was concentrated then diluted in EtOAc (200 mL) and washed with $H_2O$ (100 mL) and brine (x2, 100 mL). The isolated organic layer was dried over $Na_2SO_4$ then concentrated to dryness and purified by column chromatography (0–2% MeOH/$CH_2Cl_2$) to yield the product (361 mg, 45%). $^1$H NMR (400 MHz, MeOD-$d_4$) δ 8.06 (dd, $J$ = 7.7, 1.4 Hz, 1H), 7.87 (s, 1H), 7.54 (d, $J$ = 7.0 Hz, 2 H), 7.50 (dd, $J$ = 7.7, 1.5 Hz, 1H), 7.40 (m, 3H), 7.34 (dt, $J$ = 7.2, 1.5 Hz, 1H), 7.01 (d, $J$ = 7.7 Hz, 1H), 5.73 (s, 2H), 5.38 (s, 2H), 3.94 (s, 3H).

**6**, step b: *Methyl 2-((6-(benzyloxy)-2-(bis(tert-butoxycarbonyl)amino)-9H-purin-9-yl)methyl)benzoate*. Purine from step a (85 mg, 0.21 mmol) was diluted in MeCN (0.12 M, 5 mL) then treated with $Et_3N$ (280 µL, 1.96 mmol), $Boc_2O$ (280 mg, 1.26 mmol), and DMAP (12 mg, 0.10) and warmed to 70 °C. After 16 h the reaction mixture was concentrated to dryness then diluted in EtOAc (100 mL) and washed with $H_2O$ (50 mL) and brine (50 mL). The isolated organic layer was concentrated to dryness and purified by column chromatography (40–60% ethyl acetate/hexanes) to yield the Boc-protected product (70 mg, 57%). $^1$H NMR (400 MHz, $CDCl_3$) δ 8.16 (s, 1H), 8.05 (d, $J$ = 7.4 Hz, 1H), 7.52 (d, $J$ = 7.5 Hz, 2H), 7.36 (m, 6H), 7.23 (d, $J$ = 7.8 Hz, 1H), 5.85 (s, 2H), 5.65 (s, 2H), 3.94 (s, 3H), 1.39 (s, 18H).

**6**, step c: *tert-butyl (6-(benzyloxy)-9-(2-(hydroxymethyl)benzyl)-9H-purin-2-yl)carbamate*. Purine from step b (350 mg, 0.67 mmol) was diluted in THF (0.1 M, 7 mL) then treated with lithium aluminum hydride (77 mg, 2.03 mmol) at 0 °C. After 2 h the reaction mixture was quenched with a 10% solution of Rochelle's salt (5 mL) then diluted in EtOAc (200 mL) and passed through a pad of celite. The filtrate was washed with $H_2O$ (100 mL) and brine (100 mL). The isolated organic layer was dried over $Na_2SO_4$ and concentrated to dryness. The resulting crude residue was purified by column chromatography (1–2% MeOH/$CH_2Cl_2$) to provide the product (240 mg, 77%). $^1$H NMR (400 MHz, MeOD-$d_4$) δ 8.04 (s, 1H), 7.58 (dd, $J$ = 6.7 Hz, 2H), 7.44 (dd, $J$ = 7.3, 1.3 Hz,

1H), 7.35 (m, 3H), 7.31 (dd, $J$ = 4.8, 1.7 Hz, 1H), 7.27 (m, 1H), 7.22 (dd, $J$ = 7.5, 1.2 Hz, 1H), 5.65 (s, 2H), 5.62 (s, 2H), 4.79 (s, 2H), 1.59 (s, 9H).

**6**, steps d and e: *(E)-(((2-((6-(benzyloxy)-2-((tert-butoxycarbonyl)amino)-9H-purin-9-yl)methyl)styryl)phosphoryl)bis(oxy))bis(methylene) bis(2,2-dimethylpropanoate)*. Purine from step c (340 mg, 0.60 mmol) was diluted in $CH_2Cl_2$ (0.05 M, 12 mL) then treated with $NaHCO_3$ (252 mg, 3.0 mmol) and Dess-Martin Periodinane (334 mg, 0.80 mmol) at 23 °C. After 1 h the reaction mixture was quenched $H_2O$ (1 mL) then diluted in $H_2O$ (100 mL). The resulting mixture was extracted with $CH_2Cl_2$ (x2, 100 mL) then dried with $Na_2SO_4$ and concentrated to dryness. The isolated organic layer was concentrated to dryness and the resulting crude aldehyde was used without further purification. TetraPOM methylenediphosphonate (388 mg, 0.62 mmol) was diluted in THF (0.03 M, 15 mL) then treated with NaH (60% in mineral oil, 50 mg, 1.23 mmol) at 0 °C. After 5 min the crude aldehyde (230 mg, 0.41 mmol) in minimal amount of THF (1 mL) was added dropwise at 0 °C. After 1 h the reaction mixture was removed from the cooling batch and allowed to reach 23 °C. After 3.5 h the reaction mixture was quenched with $NH_4Cl_{(sat)}$ (5 mL) then concentrated and diluted further with EtOAc (300 mL). The mixture was washed with $NH_4Cl_{(sat)}$ (100 mL), $H_2O$ (200 mL) and brine (200 mL) and dried over $Na_2SO_4$. The isolated organic layer was concentrated to dryness and purified by column chromatography (40–60% ethyl acetate/hexanes) to provide the vinyl phosphonate (30 mg, 10% over 2 steps). $^1$H NMR (400 MHz, MeOD-$d_4$) δ 8.01 (s, 1H), 7.71 (d, $J$ = 7.2 Hz, 1H), 7.57 (m, 2H), 7.45 (m, 2H), 7.35 (m, 4H), 6.42 (d, $J$ = 17.3 Hz, 1H), 5.69 (dq, $J$ = 13.2, 2.4 Hz, 4H), 5.65 (s, 2H), 5.64 (s, 2H), 5.43 (dd, $J$ = 12.4, 5.1 Hz, 1H), 1.59 (s, 9H), 1.13 (s, 18H).

**6**, step f: *(((2-((2-((tert-butoxycarbonyl)amino)-6-oxo-1,6-dihydro-9H-purin-9-yl)methyl)phenethyl)phosphoryl)bis(oxy))bis(methylene) bis(2,2-dimethylpropanoate)*. Vinyl phosphonate from step e (30 mg, 0.03 mmol) was diluted in MeOH (0.01 M, 3 mL) then treated with 10% Pd/C (10 mg, 30% w/w) and back-filled with $H_{2(g)}$ at 23 °C. After 17 h the reaction mixture was filtered through a pad of celite then concentrated to dryness and purified by column chromatography (0–4% MeOH/$CH_2Cl_2$) to yield the product (20 mg, 78%). $^1$H NMR (400 MHz, DMSO-$d_6$) δ 10.30 (bs, 1H), 7.74 (t, $J$ = 2.3 Hz, 1H), 7.65 (dd, $J$ = 2.4, 0.5 Hz, 1H), 7.34 (dd, $J$ = 8.7, 2.1 Hz, 1H), 7.26 (m, 1H), 7.23 (dd, $J$ = 8.8, 0.5 Hz, 1H), 6.82 (d, $J$ = 8.1 Hz, 1H), 6.54 (bs, 1H), 5.64 (dq, $J$ = 13.1, 1.9 Hz, 4H), 5.25 (s, 2H), 2.95 (m, 2H), 2.24 (m, 2H), 1.17 (s, 9H).

**6**, steps g and h: *2-amino-9-(2-(2-(bis((pivaloyloxy)methoxy)phosphoryl)ethyl)benzyl)-7-((5-chlorobenzofuran-2-yl)methyl)-6-oxo-6,9-dihydro-1H-purin-7-ium (**6**)*. Purine from step f (20 mg, 0.03 mmol) was diluted in a mixture of $CH_2Cl_2$:TFA (2:1, 0.03 M, 1 mL) at 23 °C. After 2 h the reaction mixture was concentrated to dryness and the resulting crude guanine analog was used without further purification. Crude guanine analogue (17 mg, 0.03 mmol) was diluted in DMSO (0.03 M, 1 mL) then treated with 2-(bromomethyl)-5-chlorobenzofuran (24 mg, 0.10 mmol) and warmed to 50 °C. After 48 h the reaction mixture was loaded directly on to silica gel then purified by column chromatography (2–10% MeOH/$CH_2Cl_2$) and purified further by RP-HPLC Method A to yield cap analogue prodrug **6** (1 mg, 5%).

**Scheme 6.** Synthesis of **7**

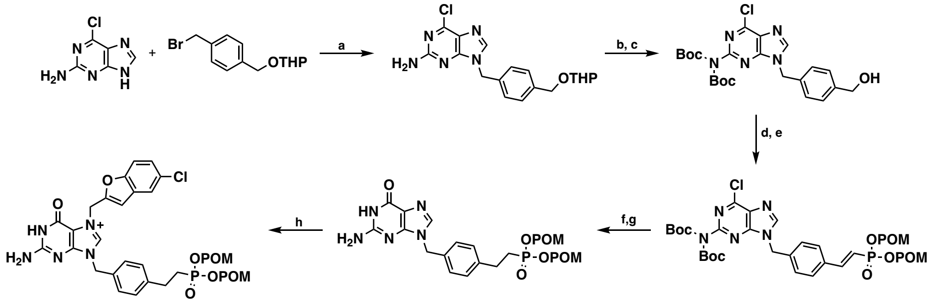


Reagents and conditions: (a) benzyl bromide, $K_2CO_3$, DMF, 80 °C; (b) $Boc_2O$, DMAP, THF, 23 °C; (c) pTSA monohydrate, MeOH, 23 °C; (d) DMP, $CH_2Cl_2$, 23 °C; (e) NaH, tetraPOM methylenediphosphonate, THF, 0 °C to 23 °C; (f) $CH_2O_2$:$H_2O$ (1:1), 40 °C; (g) 10% Pd/C, $H_2$, MeOH:$H_2O$ (4:1), 23 °C; (h) 2-(bromomethyl)-5-chlorobenzofuran, DMSO, 50 °C.

**7**, step a: *6-chloro-9-(4-(((tetrahydro-2H-pyran-2-yl)oxy)methyl)benzyl)-9H-purin-2-amine*. 2-amino-6-chloropurine (1.66 g, 9.85 mmol) was diluted in DMF (0.2 M, 50 mL) then treated with $K_2CO_3$ (4.10 g, 29.6 mmol) and 2-((4-(bromomethyl)benzyl)oxy)tetrahydro-2*H*-pyran (2.8 g, 9.85 mmol) at 80 °C. After 21 h the reaction mixture was cooled to 23 °C then concentrated to dryness and diluted in EtOAc (300 mL). The resulting organic layer was washed with $H_2O$ (200 mL) and brine (x3, 100 mL) then dried over $Na_2SO_4$ and concentrated to dryness. The resulting crude residue was purified by column chromatography (1–2% MeOH/$CH_2Cl_2$) to yield the product (1.4 g, 39%). $^1$H NMR (400 MHz, MeOD-d$_4$) δ 8.11 (s, 1H), 7.36 (q, $J$ = 4.0 Hz, 4H), 5.35 (s, 2H), 4.74 (d, $J$ = 12.2 Hz, 1H), 4.70 (t, $J$ = 4.3 Hz, 1H), 4.50 (d, $J$ = 12.2 Hz, 1H), 3.90 (m, 1H), 3.53 (m, 1H), 1.85 (m, 1H), 1.73 (m, 1H), 1.58 (m, 4H).

**7**, step b and c: *tert-butyl (tert-butoxycarbonyl)(6-chloro-9-(4-(hydroxymethyl)benzyl)-9H-purin-2-yl)carbamate*. Purine from step a (1.3 g, 3.48 mmol) was diluted in THF (0.1 M, 35 mL) then treated with $Boc_2O$ (2.28 g, 10.45 mmol) and DMAP (43 mg, 0.35 mmol) at 23 °C. After 3 d the reaction mixture was concentrated then diluted further in EtOAc (200 mL) and washed with $H_2O$ (100 mL) and brine (100 mL). The isolated organic layer was dried over $Na_2SO_4$ then concentrated to dryness to provide the crude Boc-protected product which was used without further purification. The crude product from step b (1.8 g, 3.57 mmol) was diluted in MeOH (0.1 M, 35 mL) then treated with pTSA monohydrate (64 mg, 0.34 mmol) at 23 °C. After 1h the reaction mixture was concentrated then diluted with $CH_2Cl_2$ (300 mL) and washed with $NaHCO_{3(sat)}$ (x2, 100 mL), $H_2O$ (100 mL) and brine (100 mL). The isolated organic layer was dried over $Na_2SO_4$ then concentrated to dryness and purified by column chromatography (30% ethyl acetate/hexanes) to yield the product (1.0 g, 63%). $^1$H NMR (300 MHz, $CDCl_3$) δ 8.37 (bs, 1H), 7.34 (m, 4H), 5.46 (s, 2H), 4.71 (s, 2H), 1.45 (s, 18H).

**7**, steps d and e: *(E)-(((4-((2-(bis(tert-butoxycarbonyl)amino)-6-chloro-9H-purin-9-yl)methyl)styryl)phosphoryl)bis(oxy))bis(methylene) bis(2,2-dimethylpropanoate).* Purine from step c (300 mg, 0.61 mmol) was diluted in $CH_2Cl_2$ (0.1 M, 6 mL) then treated with Dess-Martin Periodinane (336 mg, 0.79 mmol) at 23 °C. After 2 h the mixture was quenched with $Na_2S_2O_{3(sat)}$ (4 mL) then stirred for an additional 30 min and diluted further in $CH_2Cl_2$ (200 mL). The resulting mixture was washed with $H_2O$ (100 mL) and brine (100 mL) then the isolated organic layer was dried over $Na_2SO_4$ and concentrated to dryness. The crude aldehyde isolated was used without further purification. TetraPOM methylenediphosphonate (400 mg, 1.23 mmol) was diluted in THF (0.07 M, 12 mL) then treated with NaH (60% in mineral oil, 98 mg, 2.5 mmol) at 0 °C. After 5 min the reaction mixture was treated with the crude aldehyde in minimal amount of THF (~1 mL) at 0 °C. After 1 h the reaction mixture was removed from the cooling bath and allowed to stir at 23 °C. After 2 h the reaction mixture was quenched with $NH_4Cl_{(sat)}$ (5 mL) then diluted in EtOAc (200 mL) and washed with $NH_4Cl_{(sat)}$ (100 mL) and brine (100 mL). The resulting organic layer was dried over $Na_2SO_4$ then concentrated to dryness and purified by column chromatography (50–70% ethyl acetate/hexanes) to yield the vinyl phosphonate product (365 mg, 56% over 2 steps). $^1$H NMR (300 MHz, DMSO-$d_6$) δ 8.94 (s, 1H), 7.66 (d, *J* = 8.6 Hz, 2H), 7.44 (m, 1H), 7.33 (d, *J* = 11.3 Hz, 2H), 6.65 (t, *J* = 18.4 Hz, 1H), 5.66 (s, 2H), 5.61 (d, *J* = 11.8 Hz, 4H), 1.30 (s, 18H), 1.11 (s, 18 H).

**7**, steps f and g: *(((4-((2-amino-6-oxo-1,6-dihydro-9H-purin-9-yl)methyl)phenethyl)phosphoryl)bis(oxy))bis(methylene) bis(2,2-dimethylpropanoate)*. Purine from step e (300 mg, 0.37 mmol) was diluted in an aqueous mixture of formic acid (1:1, 0.05 M, 8 mL) and warmed to 40 °C. After 24 h the reaction mixture was cooled to 23 °C then concentrated to dryness and the isolated crude guanine analogue was used without further purification. The crude guanine analogue was diluted in an aqueous solution of methanol (4:1, 0.02 M, 20 mL) then treated with 10% Pd/C (60 mg, 20% w/w) and back-filled with $H_{2(g)}$ at 23 °C. After 21 h the reaction mixture was filtered through a pad of celite then concentrated to dryness and purified by column chromatography (2–8% MeOH/$CH_2Cl_2$) to yield the product (54 mg, 25% over 2 steps). $^1$H NMR (400 MHz, MeOD-$d_4$) δ 7.74 (s, 1H), 7.28 (d, *J* = 8.3 Hz, 2H), 7.24 (d, *J* = 8.3 Hz, 2H), 5.64 (d, *J* = 13.1 Hz, 4 H), 5.32 (s, 2H), 2.89 (m, 2H), 2.22 (m, 2H), 1.23 (s, 18H).

**7**, step h: *2-amino-9-(4-(2-(bis((pivaloyloxy)methoxy)phosphoryl)ethyl)benzyl)-7-((5-chlorobenzofuran-2-yl)methyl)-6-oxo-6,9-dihydro-1H-purin-7-ium (7).* Purine from step g (54 mg, 0.09 mmol) was diluted in DMSO (0.06 M, 1.6 mL) then treated with 2-(bromomethyl)-5-chlorobenzofuran (68 mg, 0.28 mmol) and warmed to 50 °C. After 48 h the reaction mixture was loaded directly on to silica gel then purified by column chromatography (2–10% MeOH/$CH_2Cl_2$) and further purified by RP-HPLC Method A to yield cap analogue prodrug **7** (28 mg, 42%).

**Scheme 7.** Synthesis of **5-PA** and **6-PA**

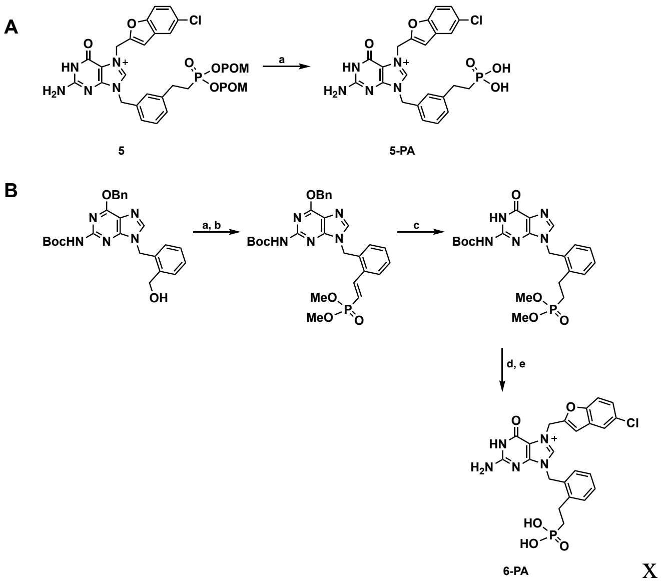


(**A**) Reagents and conditions: (a) 5N $HCl_{(aq)}$, MeOH, 45 °C; (**B**) Reagents and conditions: (a) DMP, $NaHCO_3$, $CH_2Cl_2$, 23 °C; (b) $K_2CO_3$, tetramethyl methylenediphosphonate, EtOH, 70 °C; (c) 10% Pd/C, $H_2$, MeOH, 23 °C; (d) TMSBr, MeCN, 23 °C; (e) 2-(bromomethyl)-5-chlorobenzofuran, DMSO, 50 °C.

**5-PA**, step a: *2-amino-7-((5-chlorobenzofuran-2-yl)methyl)-6-oxo-9-(3-(2-phosphonoethyl)benzyl)-6,9-dihydro-1H-purin-7-ium (**5-PA**)*. Bis-POM cap analogue **5** (40 mg, 0.053 mmol) was diluted in MeOH (0.2 M, 0.2 mL) and 5 N $HCl_{(aq)}$ (0.01 M, 5 mL) then warmed to 45 °C. After 24 h the reaction mixture was concentrated by lyophilization then purified by RP-HPLC using Method **B** to provide cap analogue **5-PA** (5.9 mg, 22%). LRMS (ESI $^{+}$) 514.1027 $[M]^{+}$.

**6-PA**, steps a and b: *tert*-butyl (*E*)-(6-(benzyloxy)-9-(2-(2-(dimethoxyphosphoryl)vinyl)benzyl)-9*H*-purin-2-yl)carbamate. Purine alcohol from scheme 5, step c (50 mg, 0.09 mmol) was diluted in $CH_2Cl_2$ (0.05 M, 2 mL) then treated with $NaHCO_3$ (37 mg, 0.45 mmol) and DMP (50 mg, 0.12 mmol) at 23 °C. After 1 h the reaction mixture was quenched with $H_2O$ (1 mL) then further diluted in $H_2O$ (100 mL) and extracted with $CH_2Cl_2$ (x2, 100 mL). The isolated organic layer was dried over $Na_2SO_4$ then concentrated and crude aldehyde was used without further purification.
The crude aldehyde was diluted in EtOH (0.02 M, 3 mL) then treated with tetramethyl methylenediphosphonate (28 mg, 0.12 mmol) and $K_2CO_3$ (33 mg, 0.24 mmol) and warmed to 70 °C. After 17 h the reaction mixture was concentrated then diluted in $CH_2Cl_2$ (100 mL) then washed with $H_2O$ (100 mL) and brine (100 mL). The isolated organic layer was dried over $Na_2SO_4$ then concentrated and purified by column chromatography (1–3% MeOH/$CH_2Cl_2$) to provide the vinyl phosphonate (42 mg, 79% over 2 steps). $^{1}$H NMR (400 MHz, $CDCl_3$) δ 10.16 (s, 1H), 7.99 (s, 1H), 7.85 (m, 1H), 7.55 (m, 4H), 7.48 (m, 1H), 7.35 (dd, *J* = 6.7,1.8 Hz, 3H), 7.32 (dt, *J* = 5.5, 1.3 Hz, 1H), 6.35 (m, 1H), 5.83 (s, 2H), 5.69 (dd, *J* = 13.7, 2.4 Hz, 2H), 5.63 (s, 2H), 3.80 (d, *J* = 10.9 Hz, 3 H), 1.56 (s, 9H)

**6-PA**, step c: *tert*-butyl (9-(2-(2-(dimethoxyphosphoryl)ethyl)benzyl)-6-oxo-6,9-dihydro-1*H*-purin-2-yl)carbamate. Vinyl phosphonate from step b (40 mg, 0.06 mmol) was diluted in MeOH (0.01 M, 6 mL) then treated with 10% Pd/C (10 mg, 20% w/w) and back-filled with $H_{2(g)}$ at 23 °C. After 17 h the reaction mixture was filtered through a pad of celite then the filtrate was concentrated and purified by column chromatography (2–6% MeOH/$CH_2Cl_2$) to provide the Boc-protected guanine analog (20 mg, 71%). $^1$H NMR (400 MHz, $CDCl_3$) δ 11.53 (s, 1H), 9.83 (s, 1H), 7.92 (s, 1H), 7.34 (d, *J* = 7.3 Hz, 1H), 7.30 (dd, *J* = 7.6, 1.3 Hz, 1H), 7.26 (d, *J* = 7.3 Hz, 1H), 7.21 (d, *J* = 7.6 Hz, 1H), 5.26 (s, 2H), 3.80 (d, *J* = 11.6 Hz, 3H), 1.55 (s, 9H).

**6-PA**, steps d and e: *2-amino-7-((5-chlorobenzofuran-2-yl)methyl)-6-oxo-9-(2-(2-phosphonoethyl)benzyl)-6,9-dihydro-1H-purin-7-ium (**6-PA**)*. Boc-protected guanine analog from step c (39 mg, 0.07 mmol) was diluted in MeCN (0.04 M, 2 mL) then treated with TMSBr (0.17 M, 0.4 mL) at 23 °C. After 20 h the reaction mixture was concentrated to dryness then diluted in $H_2O$ (10 mL) and stirred for an additional 1 h. The mixture was then concentrated *via* lyophilization to provide the guanine analog (20 mg, 87%) which was used without further purification. Guanine analog from step d (20 mg, 0.06 mmol) was diluted in DMSO (0.06 M, 1 mL) then treated with 2-(bromomethyl)-5-chlorobenzofuran (42 mg, 0.17 mmol) and warmed to 50 °C. After 24 h the reaction mixture was further diluted in $H_2O$ (50 mL) then washed with ethyl acetate (x2, 50 mL) and resulting aqueous layer was concentrated by lyophilization. The crude residue was then purified by RP-HPLC with Method B to provide the cap analogue **6-PA** (3 mg, 10%). LRMS (ESI $^+$) 514.1040 $[M]^+$.

#### D. X-ray Crystallography Data

**Table S2: Crystallography Data Collection and Refinement Statistics**

| **Data Collection** | eIF4E:**5-PA** |
|---|---|
| PDB Code | 9DON |
| SpaceGroup | $P2_1$ |
| Unit Cell a, b, c (Å) | 49.194, 74.397, 52.454 |
| Wavelength (Å) | 1.1271 |
| Resolution (Å)[1] | 2.10 (2.14-2.10) |
| Rmerge (%)[2] | 10.2 (31.6) |
| <I/sI>[3] | 14.2 (5.1) |
| Completeness (%)[4] | 98.0 (95.8) |
| Redundancy | 5.6 (5.2) |
| **Refinement** | |
| Resolution (Å) | 2.10 |
| R-Factor (%)[5] | 20.0 |
| Rfree (%)[6] | 24.7 |
| Protein atoms | 2986 |
| Water Molecules | 151 |
| Unique Reflections | 21660 |
| R.m.s.d.[7] | |
| Bonds | 0.008 |
| Angles | 0.88 |
| MolProbity Score | 1.37 |
| Clash Score[8] | 2.18 |
| Ligands | EC5121 $m^7$GDP |
| RSCC[8] | 0.089 0.86 |
| RSR[8] | 0.11 0.11 |

[1]Statistics for highest resolution bin of reflections in parentheses.

[2]$R_{sym} = \Sigma_h \Sigma_j \mid I_{hj} - \langle I_h \rangle \mid / \Sigma_h \Sigma_j I_{hj}$, where $I_{hj}$ is the intensity of observation j of reflection h and $\langle I_h \rangle$ is the mean intensity for multiply recorded reflections.

[3]Intensity signal-to-noise ratio.

[4]Completeness of the unique diffraction data.

[5]R-factor = $\Sigma_h \mid |F_o| - |F_c| \mid / \Sigma_h |F_o|$, where $F_o$ and $F_c$ are the observed and calculated structure factor amplitudes for reflection h.

[6]$R_{free}$ is calculated against a 10% random sampling of the reflections that were removed before structure refinement.

[7]Root mean square deviation of bond lengths and bond angles.

[8] wwPDB validation service.

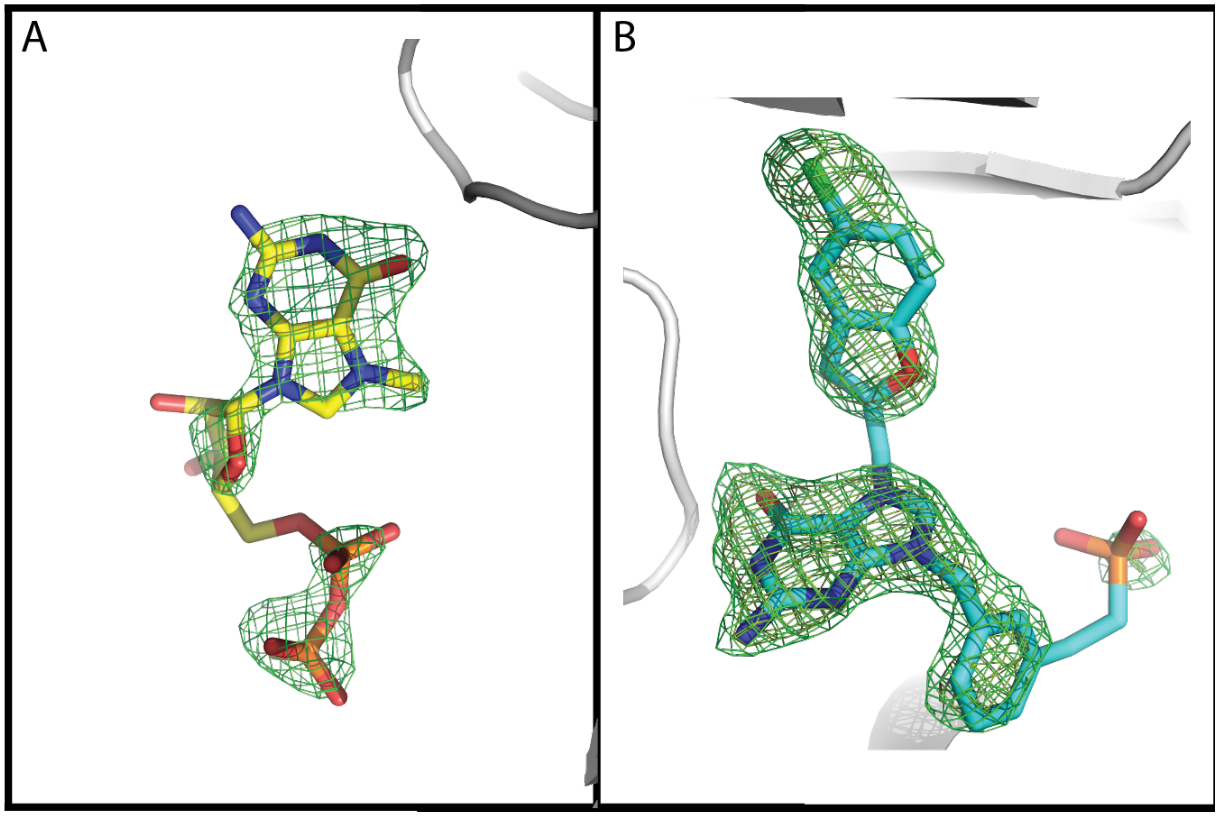


**Figure S1.** The difference electron density map resulting from the molecular replacement solution without the presence of ligands. The Fo-Fc map is represented as a grid contoured at 3.0s (light green) and 2.0s (dark green) for (A) m$^7$GDP present in chain A and (B) **5-PA** present in chain B.

#### E. References


1. Gallagher, E. E.; Song, J. M.; Menon, A.; Mishra, L. D.; Chmiel, A.; Garner, A. L., Consideration of binding kinetics in the design of stapled peptide mimics of the disordered proteins eukaryotic translation initiation factor 4E (eIF4E)-binding protein 1 (4E-BP1) and eukaryotic translation initiation factor 4G (eIF4G). *J. Med. Chem.* **2019,** *62*, 4967-4978.
2. Papadopoulos, E.; Jenni, S.; Kabha, E.; Takrouri, K. J.; Yi, T.; Salvi, N.; Luna, R. E.; Gavathiotis, E.; Mahalingam, P.; Arthanari, H.; Rodriguez-Mias, R.; Yefidoff-Freedman, R.; Aktas, B. H.; Chorev, M.; Halperin, J. A.; Wagner, G., Structure of the eukaryotic translation initiation factor eIF4E in complex with 4EGI-1 reveals an allosteric mechanism for dissociating eIF4G. *Proc. Natl. Acad. Sci., U. S. A.* **2014**, E3187-E3195.
3. Cardenas, E. L.; O'Rourke, R. L.; Menon, A.; Meagher, J.; Stuckey, J.; Garner, A. L., Design of cell-permeable inhibitirs of eukaryotic translation initiation factor 4E (eIF4E) for inhibiting aberrant cap-dependent translation in cancer. *J. Med. Chem.* **2023,** *66*, 10734-10745.
4. Otwinowski, Z.; Minor, W., *Processing of X-ray Diffraction Data Collected in Oscillation Mode*. Academic Press: New York, 1997; Vol. 276.
5. McCoy, A. J.; Grosse-Kunstleve, R. W.; Adams, P. D.; Winn, M. D.; Storoni, L. C.; Read, R. J., Phaser crystallographic software. *J. Appl. Cryst.* **2007,** *40*, 658-674.
6. Emsley, P.; Cowtan, K., Coot: model-building tools for molecular graphics. *Acta Crystallogr. D Biol. Crystallogr.* **2004,** *60*, 2126-2132.
7. Roversi, P.; Sharff, A.; Smart, O. S.; Vonrhein, C.; Womack, T. O., BUSTER version 2.11.2 Ltd., G. P., Ed. Cambridge, United Kingdom, 2011.
8. Chen, V. B.; Arendall 3rd, W. B.; Headd, J. J.; Keedy, D. A.; Immormino, R. M.; Kapral, G. J.; Murray, L. W.; Richardson, J. S.; Richardson, D. C., MolProbity: all-atom structure validation for macromolecular crystallography. *Acta Crystallogr. D Biol. Crystallogr.* **2010,** *66*, 12-21.
